# Chromosome Compartments on the Inactive X Guide TAD Formation Independently of Transcription during X-Reactivation

**DOI:** 10.1101/2020.07.02.177790

**Authors:** Moritz Bauer, Enrique Vidal, Eduard Zorita, Stefan F. Pinter, Guillaume J. Filion, Bernhard Payer

## Abstract

A hallmark of chromosome organization is the partition into transcriptionally active A and repressed B compartments and into topologically associating domains (TADs). Both structures were regarded absent from the inactive X chromosome, but to be re-established with transcriptional reactivation and chromatin opening during X-reactivation. Here, we combine a tailor-made mouse iPSC-reprogramming system and high-resolution Hi-C to produce the first time-course combining gene reactivation, chromatin opening and chromosome topology during X-reactivation. Contrary to previous observations, we uncover A/B-like compartments on the inactive X harboring multiple subcompartments. While partial X-reactivation initiates within a compartment rich in X-inactivation escapees, it then occurs rapidly along the chromosome, coinciding with acquisition of naive pluripotency, leading to downregulation of *Xist*. Importantly, we find that TAD formation precedes transcription, suggesting them to be causally independent. Instead, TADs form first within Xist-poor compartments, establishing Xist as common denominator, opposing both gene reactivation and TAD formation through separate mechanisms.

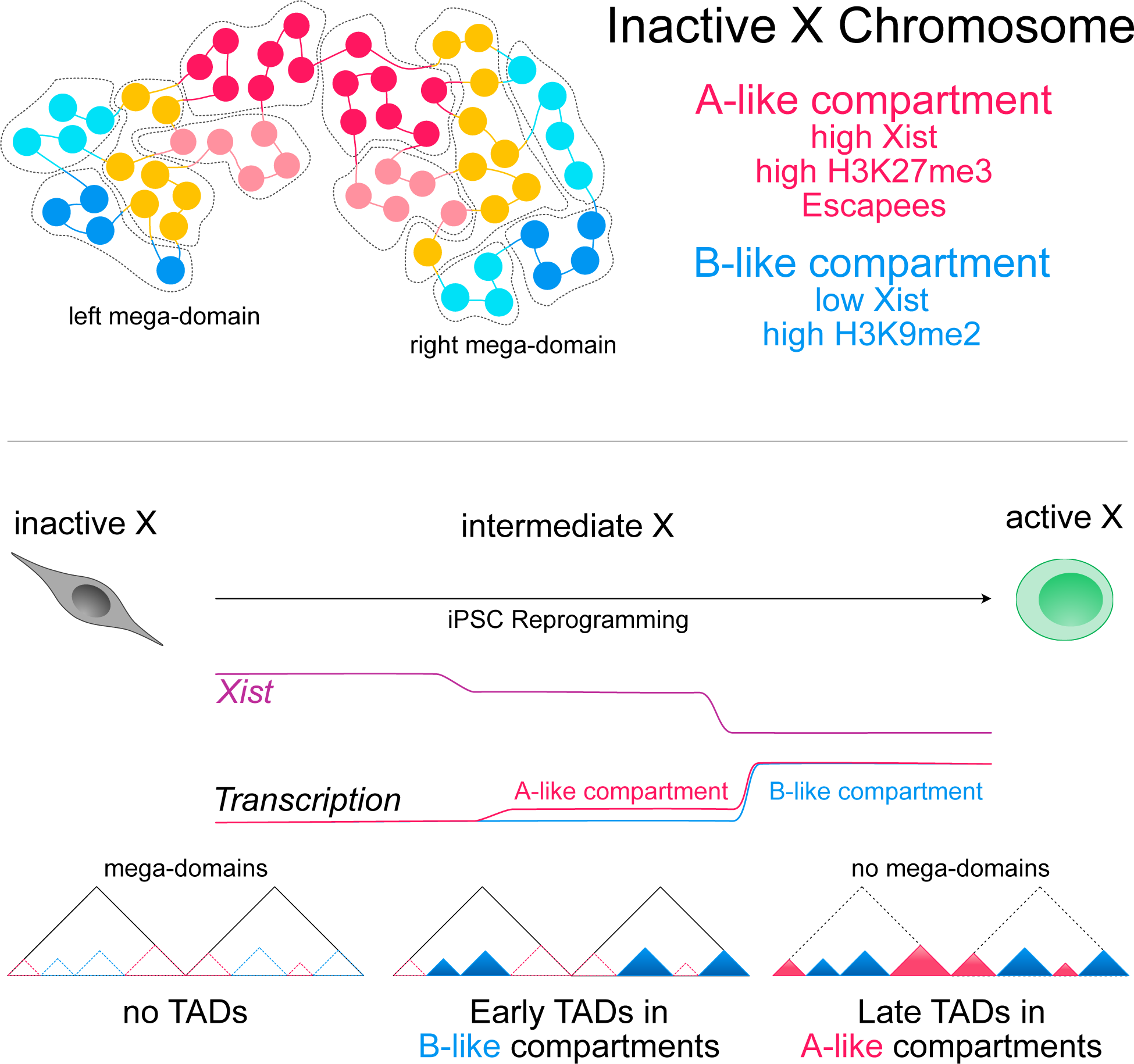
Graphical Summary.

## Introduction

To achieve gene dosage balance between males (XY) and females (XX), mammals transcriptionally inactivate one of the two X chromosomes in females during early embryonic development in a process called X-chromosome inactivation (XCI). While the active X chromosome resembles in many aspects an autosome, the inactive X (Xi) has a unique repressive configuration and chromosome conformation, which sets it apart from other chromosomes. This has established XCI as a unique model to study the formation of heterochromatin and the mechanisms of chromosome folding and of chromosome organization (Galupa and Heard, 2018; Jégu et al., 2017; Payer, 2016). Mammalian chromosomes have been shown to be organized in two separate compartments: A, corresponding to open chromatin and high RNA expression, and B, corresponding to closed chromatin and low expression (Lieberman-Aiden et al., 2009). Moreover, on a more fine-scaled level, chromosomes have been shown to be partitioned into megabase-sized local chromatin interaction domains, termed topologically associating domains (TADs), whose boundaries are enriched for the insulator binding protein CTCF and cohesin (Dixon et al., 2012; Nora et al., 2012). Intriguingly, both these levels of chromatin organization appear to be absent or attenuated on the inactive mouse X chromosome. Spatial proximity maps of the Xi obtained by Hi-C have suggested that the Xi lacks compartments (Giorgetti et al., 2016) and displays a global attenuation of TADs (Wang et al., 2018), except for regions escaping X-inactivation (Giorgetti et al., 2016; Marks et al., 2015; Minajigi et al., 2015). While the Xi has therefore been considered to be mostly “unstructured”, an exception is its unique bipartite organization of two so-called mega-domains that are separated by a tandem repeat locus, *Dxz4* (Darrow et al., 2016; Deng et al., 2015; Giorgetti et al., 2016; Rao et al., 2014). The key player in the formation of the silenced X chromosome and ultimately its unique chromosome conformation is the long non-coding RNA Xist. Xist coats the X from which it is expressed and silences the chromosome through the combined action of multiple interaction partners that set up a heterochromatic environment (Chu et al., 2015; McHugh et al., 2015; Minajigi et al., 2015). During this process, Xist repels architectural proteins like CTCF and Cohesin (Minajigi et al., 2015), thereby actively contributing to the attenuation of TADs (Wang et al., 2018) and leading to the Xi’s distinct chromosome conformation (Colognori et al., 2019; Giorgetti et al., 2016; Splinter et al., 2011). There has been intense research effort to understand the dynamics of transcriptional silencing, the mechanisms of transition to the unique structure of the Xi and the connection between the two processes (Froberg et al., 2018; Gdula et al., 2019; Wang et al., 2018, 2019), but how the process is reversed during the reactivation of the X chromosome has received attention only recently (Cantone and Fisher, 2017; Payer, 2016; Talon et al., 2019).

In mice, X-reactivation occurs twice during early development. The first round takes place at the blastocyst stage within the pluripotent epiblast of the inner cell mass (Borensztein et al., 2017; Mak et al., 2004; Okamoto et al., 2004; Payer et al., 2013). This allows the female embryo to switch from an imprinted form of X-inactivation, whereby the X inherited from the father’s sperm is inactivated, to a random form where either X can be inactivated. The second round of X - reactivation takes place in primordial germ cells during their migration and colonization of the gonads, ensuring that an active X can be transmitted to the next generation (Chuva de Sousa Lopes et al., 2008; Mallol et al., 2019; de Napoles et al., 2007; Sugimoto and Abe, 2007). However, while mechanistically, both share common features like the downregulation of *Xist* and the erasure of silencing marks like H3K27me3, their kinetics differ greatly, as X-reactivation in the blastocyst occurs within a day, while it takes several days during germ cell development.

X-reactivation can also be studied *in vitro*: induced pluripotent stem cells (iPSCs) generated from somatic cells through reprogramming have two active X chromosomes (Janiszewski et al., 2019; Maherali et al., 2007a; Pasque et al., 2014; Payer et al., 2013; Stadhouders et al., 2018), linking the X-reactivation process to de-differentiation into the naïve pluripotent stem cell state (Payer and Lee, 2014). However, the mechanisms that govern the transition have not yet been elucidated. It is unclear how the interplay of sequence, 3D-structure, chromatin status, and *trans-* acting factors affects the reactivation of X-linked genes and why X-reactivation *in vitro* during iPSC reprogramming is a slow and gradual progress as recently proposed (Janiszewski et al., 2019). In particular, it is unknown, if the dramatic topological rearrangement of the X chromosome from an inactive state with two mega-domains into an autosome-like active state consisting of A/B-compartments and TADs (Deng et al., 2015; Giorgetti et al., 2016; Minajigi et al., 2015; Rao et al., 2014) occurs before X-linked genes are reactivated, or rather follows transcription as observed during XCI (Collombet et al., 2020). This is especially relevant from a general gene regulatory point of view, as cause and effect between chromosome topology and transcriptional activity have been under debate (Hug et al., 2017; Rowley et al., 2017; Stadhouders et al., 2018). The dramatic transcriptional and structural remodeling of an entire chromosome from an OFF to an ON state makes X-reactivation a particularly attractive model system to address these important questions.

Here, we therefore set out to investigate the temporal dynamics of transcriptional reactivation, chromatin opening, and their relationship to the topological rearrangement of the inactive X in an optimized iPSC reprogramming system.

## Results

### PaX, a Novel Reporter Model System for X-Chromosome Reactivation

Previous studies on X-chromosome reactivation during iPSC reprogramming were based on mouse embryonic fibroblast (MEF) reprogramming systems (Janiszewski et al., 2019; Maherali et al., 2007b; Pasque et al., 2014; Payer et al., 2013) and have been mitigated by several limitations. First, reprogramming and X-reactivation efficiencies were low so that it has been difficult to study the process by assays that relied on a large number of cells, such as Hi -C. Second, the heterogeneity of samples was high, with cells of different reprogramming stages and degrees of X-reactivation represented in single populations.

We therefore created an optimized *in vitro* model system called PaX (Pluripotency and X chromosome reporter) designed to overcome these pitfalls (**Figure 1A**). PaX is based on a hybrid female embryonic stem cell (ESC) line (Lee and Lu, 1999; Ogawa et al., 2008), that contains one Mus musculus (X_mus_) and one Mus castaneus (X_cas_) X chromosome, allowing us to allelically distinguish the active and inactive X. Furthermore, the cell line harbors a Tsix truncation (TST) on X_mus_, forcing a biased inactivation during differentiation (Luikenhuis et al., 2001; Ogawa et al., 2008). PaX ESCs are differentiated into neural precursor cells (NPCs), to consistently obtain a large and homogeneous population of somatic cells that have undergone X-chromosome inactivation (Abranches et al., 2009). iPSC reprogramming of NPCs is initiated by the addition of doxycycline which triggers the expression of an optimized all-in-one doxycycline-inducible MKOS (c-Myc, Klf4, Oct4, Sox2) reprogramming cassette (Chantzoura et al., 2015). During the reprogramming process, the pluripotency- (P-RFP, driven by a Nanog promoter fragment) and X-reporters (X-GFP) (Wu et al., 2014) are used to isolate first, pluripotent cells poised for X-reactivation and later, cells having undergone X-reactivation. The unique features of the PaX system enabled us to isolate pure populations of cells at different stages of X-reactivation allowing us to chart a roadmap of the X-reactivation process with unprecedented time resolution (**Figure 1A**).

**Figure 1.**
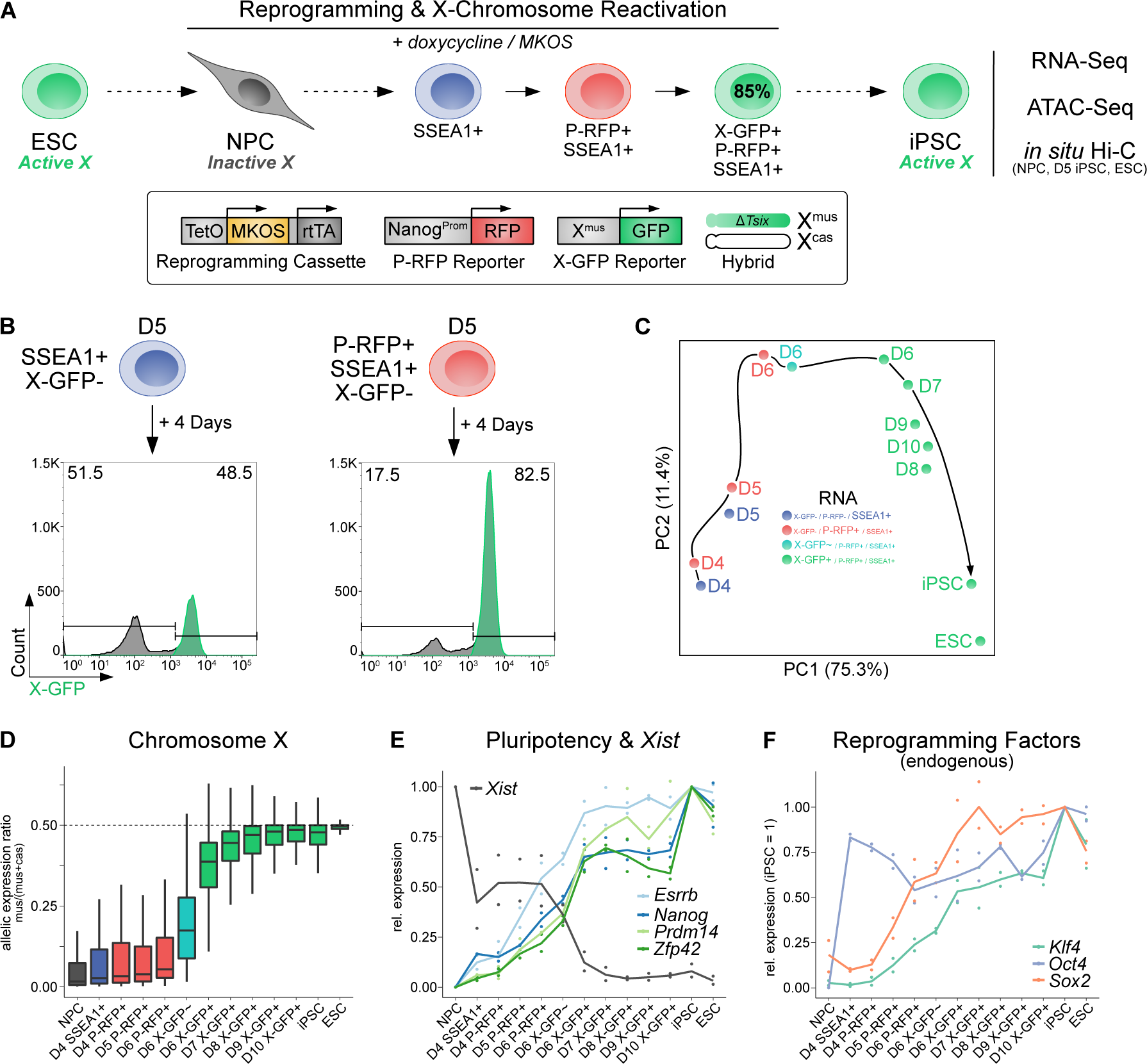
A Novel Reprogramming System to Efficiently Trace X-Chromosome Reactivation. (A) Schematic representation of the PaX reprogramming system. (B) X-reactivation efficiency of indicated reprogramming intermediates isolated on day 5 and then reprogrammed for an additional 4 days. Shown are representative histograms gated on SSEA-1+ cells. Numbers indicate the percentages of X-GFP+ and X-GFP-cells. (C) PCA of gene expression dynamics during reprogramming (n = 12,318 genes). Black arrow, hypothetical trajectory. (D) Allelic expression ratio (mus/(mus+cas)) of protein-coding genes expressed from chromosome X (n = 335). For biallelic expression, ratio = 0.5. (E) Average gene expression kinetics of naive pluripotency genes *Esrrb*, *Nanog*, *Prdm14*, and *Zfp42* (relative to the levels in iPSC) and *Xist* (relative to the levels in NPC) during reprogramming (n = 2). (F) Average endogenous gene expression kinetics of the reprogramming factor genes *Klf4*, *Oct4*, and *Sox2* during reprogramming (n = 2, relative to the levels in iPSC). Endogenous expression assessed via the genes’ 3’-UTR.

First, we assessed the kinetics of reprogramming markers with our PaX reporter cell line. After initiation of reprogramming, our cell line first upregulates the pluripotency marker stage-specific embryonic antigen 1 (SSEA1) with subsequently around 15-25% of SSEA1+ cells becoming P-RFP+ (**Figure S1A**), of which up to 85% reactivate X-GFP (**Figure S1B**). To test if P-RFP+/X-GFP-cells therefore represented a pluripotent population primed for X-reactivation, we isolated SSEA1+/P-RFP-/X-GFP- and SSEA1+/P-RFP+/X-GFP-cells on day 5 by fluorescence-activated cell sorting (FACS) and continued reprogramming. Analysis 4 days later showed that while only half of the SSEA1+/P-RFP-cells were able to reactivate X-GFP, the number rose to 80% for SSEA1+/P-RFP+ (**Figure 1B**). We conclude that our PaX system enables us to separate homogeneous cell populations, which is a prerequisite for a faithful kinetic analysis of the X - reactivation process.

Utilizing this specialized reprogramming system, we set out to obtain a high-resolution map of X-reactivation in relation to the iPSC-reprogramming process. We performed differentiation of PaX ESCs into NPCs followed by reprogramming to iPSCs and sorted intermediate reprogramming stages in 24 h intervals. On these subpopulations, we performed allele-specific RNA-Seq and ATAC-Seq (Assay for Transposase-Accessible Chromatin with High Throughput Sequencing) to reveal gene reactivation and chromatin opening kinetics, respectively. We also performed *in situ* Hi-C at three key stages (NPCs, D5 and ESCs) to get the first overview of the structural changes during X-reactivation at high resolution.

To define the trajectory towards X-reactivation, we performed principal component analysis (PCA) of the RNA-Seq data. As day 4 (D4) samples clustered far away from NPCs (**Figure S1C**), we repeated this analysis excluding NPCs to improve the resolution along the reprogramming time course (**Figure 1C**). This revealed a trajectory of reprogramming and X-reactivation with states that would have been merged otherwise. Next, to determine X chromosome-wide gene reactivation kinetics along this trajectory, we assessed the allelic expression ratio between the inactive X_mus_ and the active X_cas_, for 335 genes which had sufficient allelic information and expression (*see methods*). Whereas D6 P-RFP+/XGFP-cells still portrayed inactivation levels similar to NPCs, we found a clear switch in D6 P-RFP+/X-GFP+ cells, displaying an allelic ratio close to iPSCs (**Figure 1D**), showing reactivation of the inactive X_mus_ in this population. In contrast, genes on chromosome 13 maintained a consistent biallelic expression throughout reprogramming (**Figure S1D**), confirming that these allelic changes were specific to the inactive X_mus_.

In parallel, we observed the characteristic sequential activation of endogenous pluripotency factors (**Figures 1E and 1F**), a key event during iPSC reprogramming (Di Stefano et al., 2014; Polo et al., 2012; Stadhouders et al., 2018; Stadtfeld et al., 2008). First, we could detect high endogenous expression levels of *Oct4* on D4, at which point we also observed a sharp drop in expression markers of neural precursor cells *Blbp*, *Nestin*, *Pax6*, and *Sox1*, showing the rapid extinction of the somatic gene expression signature (**Figure S1E**). On D5, we observed activation of *Sox2* and *Esrrb* and on D6 upregulation of naive pluripotency factors *Klf4*, *Nanog*, *Prdm14,* and *Zfp42/Rex1,* coinciding with downregulation of the X-inactivation master regulator *Xist* (**Figure 1E**). This is consistent with *Xist* downregulation during iPSC reprogramming being dependent on binding of both core and naive pluripotency factors along their binding hubs at the *X-chromosome inactivation center* (*Xic*) (Gontan et al., 2012; Navarro et al., 2008, 2011; Payer and Lee, 2014; Payer et al., 2013). In conclusion, the unique properties of the PaX system revealed that X-reactivation during iPSC reprogramming occurs in a switch-like synchronous fashion, and is tightly linked to the establishment of the naive pluripotency program and the downregulation of *Xist*. Thereby it faithfully mirrors the rapid X-reactivation kinetics observed in mouse blastocysts *in vivo* (Borensztein et al., 2017).

### An Underlying A/B-like Compartmentalization Persists on the Inactive X Chromosome

Previous studies have shown that the active and inactive X chromosome have strikingly different 3D conformations (Bansal et al., 2019; Deng et al., 2015; Giorgetti et al., 2016; Minajigi et al., 2015; Nora et al., 2012; Rao et al., 2014; Splinter et al., 2011). In particular, while the active mouse X chromosome has an autosome-like structure, exhibiting active A and inactive B compartments and topologically associating domains (TADs), the inactive X chromoso me is thought to lack compartmentalization and to solely exhibit TADs of attenuated strength (Wang et al., 2018). Moreover, the inactive X consists of two mega-domains divided by the boundary element *Dxz4* with TAD structures only around genes that escape the X-inactivation process (Giorgetti et al., 2016). The so-called “unstructured” state of the inactive X is shaped by xist RNA, which repels CTCF and cohesins, thereby causing the loss/attenuation of TADs (Minajigi et al., 2015) and by SMCHD1, which is responsible for the merging of compartments on the inactive X chromosome (Wang et al., 2018). We thus obtained the contact map of the X chromosome using Hi-C to determine how structural remodeling ties in with chromatin opening and transcriptional reactivation during X-reactivation.

We performed *in situ* Hi-C (Rao et al., 2014; Stadhouders et al., 2018) on NPCs, D5 P-RFP+/X-GFP-pre-iPSCs, and ESCs (**Figures 2A** **and S2A**), *i.e.*, at the endpoints and immediately before the reactivation of gene expression. Furthermore, we FACS-sorted G1 cells based on DNA content (**Figure S2B**), which reduces cell cycle-induced variability and recovers a greater proportion of long-range *cis* contacts of samples (Bonev et al., 2017). Replicates were highly correlated (**Figure S2C**) and reached a considerably higher resolution than comparable studies assessing the murine inactive X using *in situ* Hi-C (**Figure S2D**) (Wang et al., 2018, 2019). Visual inspection of the Hi-C matrices revealed the characteristic presence of the two mega-domains on the inactive X_mus_, not only in NPCs but also in D5 P-RFP+ cells (**Figure 2A**), showing that the mega-domain structure remains for at least 5 days into the reprogramming process.

**Figure 2.**
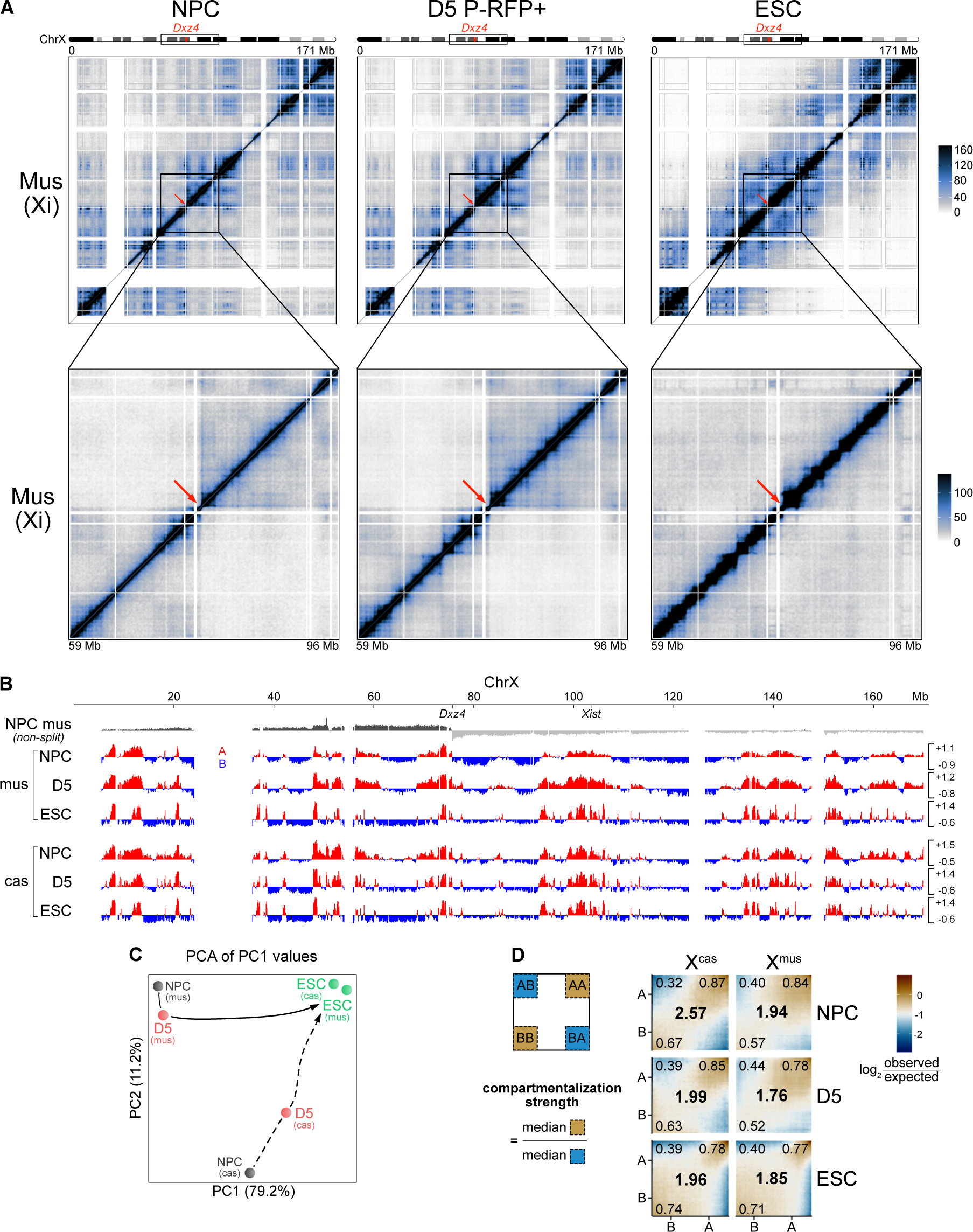
The Inactive X Chromosome Exhibits A/B-Like Compartmentalization. (A) Allele-specific Hi-C maps of chromosome X_mus_ at the inactive state in NPCs (left), intermediate state during reprogramming in D5 P-RFP+ cells (middle) and in the active state in ESCs (right). Top: Entire chromosome is shown at 200-kb resolution. Bottom: Zoom-in of the mega-domain boundary is shown at 100-kb resolution. Scale is shown in mega-bases (Mb). The mega-domain boundary *Dxz4* is indicated by a red arrow. White-shaded areas, unmappable regions. (B) A/B compartments of chromosome X at 100-kb resolution obtained with principal component analysis of matrices split at the *Dxz4* mega-domain boundary. Positive PC1 values represent A-like compartments (red); negative PC1 values represent B-like compartments (blue). Top: when matrices are not split at the *Dxz4* mega-domain boundary, then the PC1 corresponds to the two mega-domains for the inactive X chromosome. (C) PCA of PC1 values to compare A/B-like compartmentalization of the X_mus_ and X_cas_ at different stages. (n = 1,406 bins). Black arrow, hypothetical trajectory. (D) Saddle plots showing the interactions within (AA, BB) and between (AB, BA) compartments (small numbers in the corners) of chromosome X. Data are presented as the log_2_ ratio of observed versus expected aggregated contacts between bins of discretized eigenvalues (50 categories, bin size = 100 kb). Overall compartmentalization strengths for X_mus_ and X_cas_ at different stages are shown as large numbers in the center.

Unexpectedly, we could observe the distinct checkerboard pattern associated with genomic compartment structures (Lieberman-Aiden et al., 2009) on the Xi in NPCs, which prompted us to further investigate the possible compartmentalization of the Xi. We applied PCA on our Hi -C data and initially found that the first eigenvector (‘PC1 values’) captured the two mega-domains of the Xi (**Figure 2B** top) as previously reported (Giorgetti et al., 2016). However, we reasoned that the dominant mega-domain boundary at *Dxz4* may obscure an underlying compartment structure, and therefore repeated this analysis on each mega-domain separately. Strikingly, this revealed an underlying compartment structure on the inactive X_mus_ in NPCs (**Figure 2B**). We identified ∼75 A/B-like compartments on the Xi, which visually resembled the ∼90 A/B compartments on the Xa in NPCs, which themselves were distinct from the ∼120 A/B compartments of the two active X chromosomes in ESCs. Furthermore, PCA showed that the compartment structure of the Xi remained stable on D5 when on the contrary, the Xa structure had already been partially remodeled (**Figure 2C**), in accordance with the genome-wide restructuring of A/B-compartments during the transition from a somatic to a pluripotent state (Stadhouders et al., 2018). This suggests that the presence of Xist and its associated chromatin state delay changes of the Xi compartment structure until Xist becomes fully downregulated during reactivation (**Figure 1E**).

Next, we wanted to address if compartmentalization of the Xi was observable in our datasets due to the increased Hi-C resolution and G1-sorting in our data set or if it could have been observed in other datasets as well. We re-analyzed allele-specific *in situ* Hi-C data from NPCs (Wang et al., 2018) and mouse embryonic fibroblasts (MEFs) (Wang et al., 2019), and found that both exhibited compartmentalization of the inactive X (**Figure S2E**). We noted however that the data seemed visually noisier, suggesting that both high resolution and G1 sorting contributed to revealing the compartment structure of the Xi. In summary, this indicates that the A/B-like compartmentalization is not merely restricted to our differentiation system or cell type, but is a general property of the inactive mouse X chromosome.

While we were able to unveil an underlying compartment structure on the Xi, it remained possible that the A/B-like compartments identified within the dominant mega-domain structures were similarly attenuated as TADs (Wang et al., 2018) on the Xi and were hence diminished in their ability to spatially separate the chromosome. We, therefore, measured the overall interaction strengths within and between A-like and B-like compartments and visualized these differences in compartmentalization using saddle plots (**Figure 2D**). We further computed the compartmentalization strength as a means to assess the degree of A/B spatial separation (Stadhouders et al., 2018). While we observed that compartmentalization strength of the active X_cas_ in NPCs was higher than that of the inactive X_mus_, we found overall that the compartmentalization strength of the inactive X_mus_ in NPCs is comparable or even higher than on D5 or in ESCs, therefore confirming our observation of the compartmentalization of the inactive X.

In summary, we have unveiled a previously overlooked A/B-like compartment structure on the inactive X chromosome, which is stably maintained during reprogramming until the X becomes reactivated.

### The Inactive X Structure Consists of Clusters and Subcompartments with Distinct Epigenetic Properties

In front of our unexpected discovery of an A/B-like compartment structure on the inactive X, we considered the possibility that further, more fine-grained levels of structural organization might exist. High-resolution Hi-C maps have enabled the discovery of an additional layer of organization that splits A/B compartments into 5 subcompartments (Rao et al., 2014). These were not only shown to exhibit distinct interaction patterns but additionally displa yed specific patterns of chromatin modifications (Rao et al., 2014), nuclear positioning (Chen et al., 2018; Quinodoz et al., 2018), and chromatin interaction stability (Belaghzal et al., 2019).

To investigate if such a sub-compartmentalization exists on the inactive X chromosome as well, we utilized a previously reported approach (Lucic et al., 2019) to segment our allelically-resolved Hi-C matrices into spatial clusters based on their intra-chromosomal interaction pattern (**Figure 3A**). We used matrices of the inactive X_mus_ of D5 P-RFP+ pre-iPSCs to define spatial clusters, as they capture the specific transitory stage structure of reprogramming (see below). Our spatial segmentation yielded 12 clusters, 5 on the left and 7 on the right mega-domain (**Figure 3B**), with an average domain size of 317 kb. Moreover, interaction patterns of these clusters were highly similar in NPCs, on both the inactive X_mus_ and the active X_cas_ (**Figure S3A**), in line with their overall resemblance in A/B-like compartment structure (**Figure 2B**). We then consolidated these clusters into 5 subcompartments based on their mutual interaction patterns (**Figure 3A**) and their PC1 values in NPC X_mus_ (**Figure 3C**), revealing two A-like subcompartments (A1 and A2), one intermediate subcompartment we called AB-like and two B-like sub-compartments (B1 and B2) (**Figure 3D**). Similarly to subcompartments identified globally in human cells (Rao et al., 2014), we found subcompartments A1 and A2 to be the most gene-rich on the inactive mouse X chromosome (**Figure 3E**). Accordingly, we also found A-like subcompartments to be enriched in SINE repeats (**Figure S3B**), whereas LINE1 elements were preferentially enriched in B-like subcompartments (**Figure S3C**) (Richardson et al., 2015).

**Figure 3.**
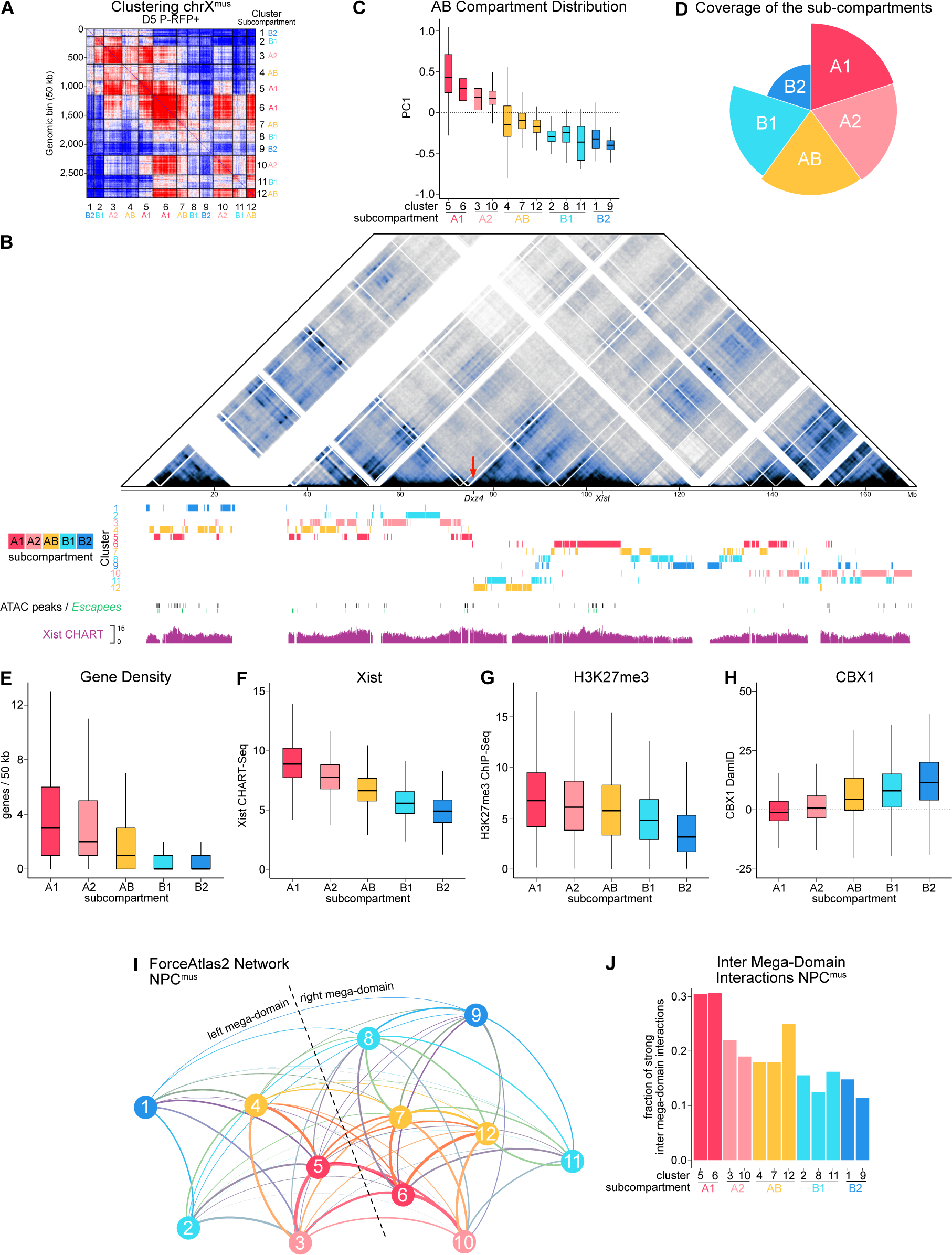
Subcompartmentalization of the Inactive X Chromosome. (A) Identification of spatial clusters (numbers 1 to 12 on the axes) and their associated subcompartments (text in color next to cluster labels) on the inactive X using *k*-means clustering on a balanced matrix of chromosome X_mus_ D5 P-RFP+ at 50-kb resolution. Red areas interact more while blue areas interact less. (B) Allele-specific Hi-C map of chromosome X_mus_ in NPCs at 100-kb resolution. Scale is shown in mega-bases (Mb). Mega-domain boundary *Dxz4* is indicated by a red arrow. White-shaded areas, unmappable regions. Position of spatial clusters is shown below. Position of ATAC peaks in NPC X_mus_ is shown in black, genes escaping X-inactivation in NPCs are shown in green. Xist RNA binding pattern in NPCs (CHART-Seq, composite scaled tracks) taken from (Wang et al., 2018). (C) Distribution of PC1 values in NPC X_mus_ of the spatial clusters. (D) Polar chart showing the coverage of the sub-compartments on chromosome X (fraction of linear sequence occupied by each subcompartment). (E) Gene density of subcompartments as number of genes per 50 kb bin. (F) Xist RNA enrichment of subcompartments in NPCs (composite scaled data). CHART-Seq data from (Wang et al., 2018). (G) H3K27me3 enrichment of subcompartments in NPC_mus_. ChIP-Seq data from (Wang et al., 2018). (H) CBX1 enrichment of subcompartments in NPC_mus_. DamID data from (Wang et al., 2018). (I) Network of spatial clusters on chromosome X_mus_ in NPCs obtained by applying the ForceAtlas2 algorithm to Hi-C interaction patterns of spatial clusters. Each cluster represents a single node of the network. Line-width correlates with interaction strength. (J) Inter-mega-domain interactions of clusters (across the mega-domain boundary) in NPC_mus_.

Because Xist initially targets gene-rich regions during XCI (Simon et al., 2013), we asked how such preferential Xist enrichment would be reflected across subcompartments on the inactive X in NPCs. To this end, we integrated published Xist CHART-Seq and ChIP-Seq data sets obtained from NPCs (Wang et al., 2018) with our compartment data and indeed found differential enrichment of Xist RNA along the subcompartments, with the highest levels of Xist detected in subcompartment A1, which harbors the *Xist* locus itself, and then gradually decreasing in the other compartments towards reaching the lowest levels in B2 (**Figures 3B and 3F**). Considering this differential Xist enrichment, we wondered if this would lead to a distinct epigenetic makeup of subcompartments. In line with polycomb-recruitment to the inactive X and Xist RNA spreading being interdependent (Colognori et al., 2019; Napoles et al., 2004; Plath et al., 2003; Silva et al., 2003), we found H3K27me3 to be enriched in A-like subcompartments (**Figure 3G**). On the contrary, H3K9me2-associated protein CBX1 was enriched in B-like subcompartments (**Figure 3H**). Moreover, consistent with previous work demonstrating repulsion of the architectural proteins CTCF and the cohesin RAD21 by Xist from the inactive X (Minajigi et al., 2015), we observed a correlation of Xist enrichment with the reduction of these factors (**Figures S3D and S3E**).

An intrinsic property of chromosomal compartments is the spatial segregation of distinct chromatin states (Lieberman-Aiden et al., 2009). Having unveiled subcompartments on the inactive X with specific epigenetic signatures, we therefore wondered how this may shape the overall structure of the X chromosome. We reasoned that utilizing the intra-chromosomal interaction pattern of these clusters would allow us to deduce their spatial relation with each other. We used ForceAtlas2 (Jacomy et al., 2014), to construct a force-directed network using either NPC or D5 P-RFP+ X_mus_ matrices, where each cluster has been consolidated into a single node (**Figures 3I** **and S3F**). This analysis revealed a spatial organization with clusters of B-like subcompartments occupying the exterior of the chromosome, with a generally low degree of connectivity. Clusters of subcompartment AB were found to be situated at the interface between A- and B-like clusters, with A-like clusters occupying the center of the network. Moreover, A1-like clusters 5 in the left and 6 in the right mega-domain appear to reside in a spatial location where the two mega-domains come closest to each other. To confirm this observation, we quantified the degree of 3D-interactions bridging the mega-domain boundary. Indeed, we found A1-like clusters (5 and 6) to exhibit the highest degree of inter mega-domain interactions (**Figures 3J** **and S3G**). It is of interest to note that the *Xist* locus resides within cluster 6 (**Figure 3B**). Therefore the close 3D-proximity between clusters 5 and 6 may facilitate the efficient spreading of Xist RNA within the gene-rich A1 compartment, which occurs first during X-inactivation (Engreitz et al., 2013; Simon et al., 2013). Xist RNA and the distinct epigenetic status of the A-like and B-like domains could subsequently contribute to the maintenance of the stable underlying compartment structure on the Xi as suggested previously (Wang et al., 2019), in contrast to the variable cell-type-specific A/B-compartments on the Xa (**Figure 2C**).

In summary, we have unveiled a subcompartment structure on the inactive X that is characterized by preferential binding of Xist RNA to gene-rich A-like compartments. Moreover, although being separated by the mega-domain boundary, gene-rich compartments are in close spatial proximity, suggesting that the 3D-structure of the X chromosome could provide a scaffold for efficient Xist-mediated gene silencing and dosage compensation.

### Chromatin Opening Emerges from Previously Open Escapee Regions and Defines Regions of Early X-Reactivation

Our identification of a fine-grained subcompartment structure on the inactive X prompted us to ask how this might impact gene expression and chromatin accessibility. Specifically, we wanted to know if the dynamics of X-reactivation might be influenced by these structural features.

When we assessed the allelic expression of the 12 spatial clusters, it became apparent that genes escaping X-inactivation in NPCs (**Figure S4A**) almost exclusively resided in subcompartment A1 (**Figure 4A**), with up to 30% of genes in cluster 5 being escapees. Moreover, while on the active X_cas_ chromatin accessibility was generally higher in the A-type compartment compared to the rest, we found on the inactive X_mus_ chromatin accessibility specifically of cluster 5 to be at much higher levels when compared to the rest of the clusters (**Figure 4B**).

**Figure 4.**
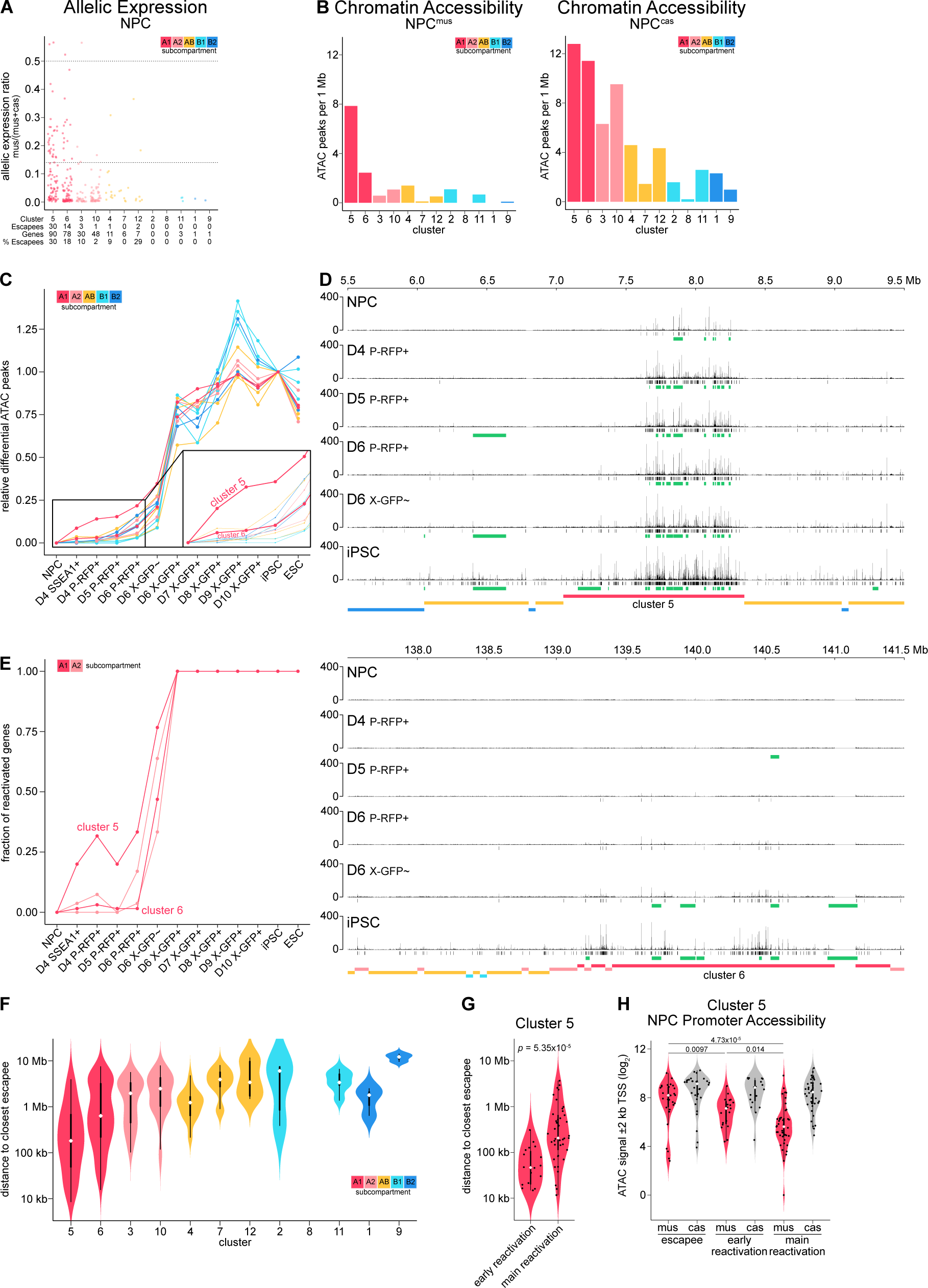
Initiation of Chromatin Opening and Gene Expression from a Distinct 3D Cluster. (A) Allelic expression ratio (= mus/(mus+cas)) of X-linked genes in spatial clusters. Cutoff >0.14 defines escapees (**Figure S4A**). For biallelic expression, ratio = 0.5. Only protein-coding genes with sufficient allelic information and expression for chromosome X_cas_ are counted (*see methods*). (B) Chromatin accessibility of each spatial cluster in NPCs shown as number of ATAC peaks per 1 Mb. (C) Dynamics of chromatin opening of spatial clusters. Only new peaks differential from NPCs were used. Relative differential ATAC peaks were then obtained by dividing the sum of peaks of each cluster at a given time point, by the sum of peaks in iPSC. Therefore NPCs will have a value of 0 and iPSCs a value of 1. Zoom-in shows early chromatin opening from NPCs until D6 P-RFP+. (D) ATAC-Seq profiles of chromatin opening at two representative X-linked regions of 4 Mb. Position of ATAC peaks is shown in black (except for NPCs, differential new peaks compared to NPCs are shown). Genes either escaping X-inactivation in NPCs or being reactivated based on RNA expression are shown in green. Position of spatial clusters is shown at the bottom. (E) Dynamics of gene reactivation of gene-rich A-like clusters. Fractions of reactivated genes per cluster are shown. 0, no reactivated gene. 1, all genes reactivated. Threshold for gene reactivation, allelic expression ratio >0.14. (F) Violin plots showing the linear distance of genes to the closest escapee. (G) Violin plots showing the linear distance to the closest escapee for genes of cluster 5 reactivating early, at D4 P-RFP+, compared to genes reactivating after that (“main reactivation”). P value calculaled by Wilcoxon rank-sum test. (H) Violin plots showing the promoter accessibility of genes of cluster 5. ATAC signal in a window of +-2 kb around the transcriptional start site (TSS) was summed. The p values are calculated by Wilcoxon rank-sum test.

These observations motivated us to ask if this would advance the timing of chromatin opening and gene reactivation in cluster 5. Indeed, when we assessed the dynamics of chromatin opening of the Xi (X_mus_) during reprogramming, we found early chromatin opening at days 4 to 6 to specifically occur in cluster 5 (**Figures 4C and 4D**). However, early chromatin opening of cluster 5 was restricted to around 25% of iPSC levels, with the most significant opening at time point D6 X-GFP+, like all the other clusters (**Figure 4C**). Similarly, when we assessed gene reactivation dynamics of the gene-rich A-like clusters (**Figures 4E** **and S4B**), we specifically observed about one-quarter of cluster 5 genes to reactivate early. Analogous to chromatin opening, early gene reactivation was restricted to around 25% of iPSC levels (**Figure S4C**).

Why do early chromatin opening and early gene reactivation happen almost exclusively in cluster 5? First, we hypothesized that higher absolute expression levels in NPCs might aid earlier reactivation. However, when we compared the expression levels of early and main reactivating genes on X_mus_ in NPCs (**Figure S4D**), we found no significant differences. Next, we asked if early reactivating genes might be bound by a distinct set of transcription factors expressed early during reprogramming. We set out to identify enriched transcription factor binding motifs using the MEME suite (McLeay and Bailey, 2010), comparing differential ATAC-Seq peaks of early reactivating genes at D4 P-RFP+, to main reactivating genes at D6-RFP+ (**Figure S4F**). However, we could not detect any significantly enriched differential motifs (**Figure S4F**), suggesting that binding of specific transcription factors is unlikely to be the main driver in directing early gene reactivation.

As we showed that cluster 5 harbors the highest percentage of escapee genes, we considered that close distance to escapees, and therefore also close vicinity to open regions, might facilitate early reactivation as shown previously (Janiszewski et al., 2019). Indeed, we found that in general, genes in cluster 5 were in closest proximity to escapees (**Figure 4F**). Compellingly, when we specifically determined the distance to escapees for early and main reactivating genes within cluster 5, we found that early genes were in significantly closer proximity to escapees than the rest of genes (**Figure 4G**). Additionally, we observed a significant enrichment of SINE elements near promoters of escapees and early reactivated genes in cluster 5 (**Figure S4G**), a property previously described for escapees and genes prone for reactivation after Xist-depletion (Hong et al., 2017; Loda et al., 2017). Moreover, while we determined that this did not affect expression levels of these genes in NPCs, where they were still transcriptionally inactive ( **Figure S4D**), we did find promoters of early reactivating genes to already be more accessible in NPCs (**Figure 4H**).

This is in line with our observation, that chromatin opening at gene promoters precedes transcription during the initiation of X-chromosome reactivation (**Figures 54H** **and I).**

In summary, partial reactivation of genes early on during reprogramming is confined to a distinct spatial cluster that is characterized by a high number of genes escaping X-inactivation.

### Remodeling of the *X-Inactivation Center* Leads to *Xist* Downregulation During Reprogramming

A critical event during X-reactivation is the downregulation of *Xist* (Pasque et al., 2014; Payer et al., 2013), which coincides with the upregulation of the naive pluripotency network (**Figures 1E and 1F**) and with the main occurrence of chromatin opening and reactivation of X-linked genes (**Figures 1D and 4C**). In order to derive mechanistic insights into the *Xist* downregulation process during reprogramming, we examined the changes of *Xist* and its known regulators at the X-inactivation center (*Xic)* (**Figures 5 and S5**).

**Figure 5.**
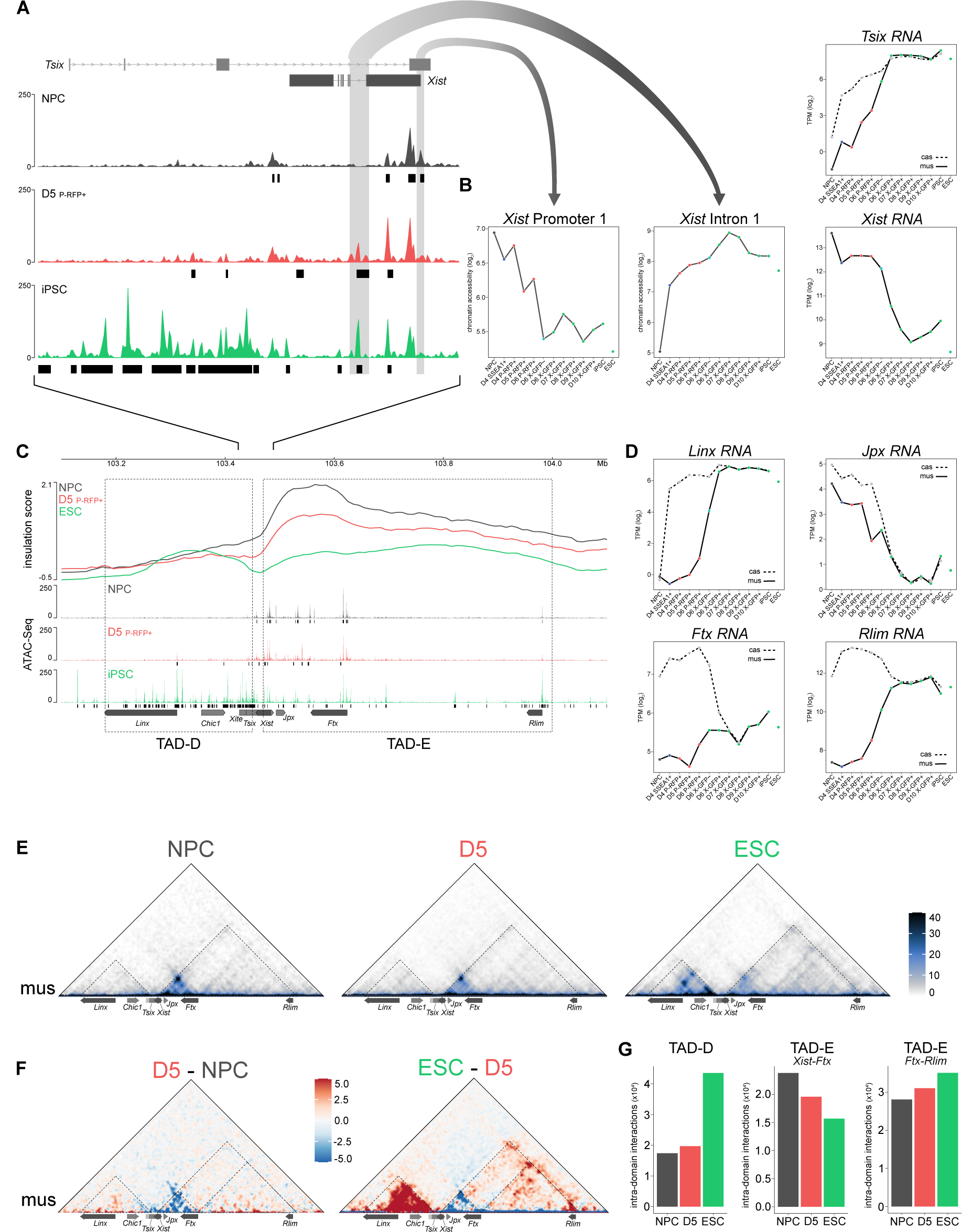
Remodeling of the X-inactivation Centre Leading to *Xist* Downregulation. (A) ATAC-Seq profiles of chromatin opening at a region encompassing the *Tsix* and *Xist* genes (mm10; 103,416,500 bp - 103,490,000 bp). Position of ATAC peaks is shown in black (except for NPCs, differential peaks compared to NPCs are shown). (B) Chromatin accessibility on X_mus_ at *Xist* promoter 1 (mm10; 103,482,600 bp - 103,483,800 bp) and *Xist* intron 1 (mm10; 103,470,900 bp - 103,474,200 bp) as depicted in (**A**). RNA expression of *Tsix* (X_mus_ and X_cas_) and *Xist* (X_mus_ only). (C) Insulation and chromatin opening at a region encompassing *Tsix* TAD-D (mm10; 103.18 Mb - 103.45 Mb) and *Xist* TAD-E (mm10; 103.47 Mb - 104.0 Mb). Top, insulation score at 10-kb resolution is shown. Bottom, ATAC-Seq profiles and ATAC-peaks in black (except for NPCs, differential peaks compared to NPCs are shown. Only genes with implicated roles in X-inactivation or X-reactivation are shown. (D) RNA expression of *Linx*, *Jpx*, *Ftx*, and *Rlim* (X_mus_ and X_cas_). (E) Allele-specific Hi-C maps of chromosome X_mus_ at 10-kb resolution at a region encompassing TAD-D (left) and TAD-E (right). Dotted lines show TADs. (F) Differential allele-specific Hi-C maps of chromosome X_mus_ at 10-kb resolution at a region encompassing TAD-D (left) and TAD-E (right). Dotted lines show TADs and additionally separate TAD-E in two regions at the TSS of *Ftx* (mm10; 103.62 Mb) for quantification in (**G**). (G) Sum of intra-domain interactions are shown. TAD-E was separated in two regions at the TSS of *Ftx*.

We first focussed on the chromatin status of the *Xist* locus itself (**Figures 5A, 5B, and S5A**). Specifically, we observed a gradual reduction in accessibility at the main *Xist* promoter 1, which preceded the full downregulation of Xist RNA during reprogramming. Following the opposite trend, we saw a gain in accessibility at *Xist* intron 1, a known binding hub for pluripotency factors such as OCT4, SOX2, NANOG, and PRDM14 (Gontan et al., 2012; Navarro et al., 2008, 2011; Payer and Lee, 2014; Payer et al., 2013). Like *Xist* RNA downregulation, gain in accessibility at *Xist* intron 1 took place in two phases: The first step occurred around D4, with a strong accessibility gain at *Xist* intron 1 and a two-fold reduction in Xist RNA levels, presumably due to expression of the reprogramming cassette, as well as the rapid reactivation of endogenous *Oct4* expression (**Figure 1F**). The second step occurred on D6 and involved another upward shift in *Xist* intron 1 accessibility and a complete downregulation of Xist RNA (**Figures 5B** **and S5B**). This coincided with a sharp increase in expression of naive pluripotency factor genes, such as *Nanog, Zfp42/Rex1* and *Prdm14* (**Figures 1E and 1F**) in line with their known role in repressing *Xist* in ESCs or during iPSC reprogramming (Gontan et al., 2012; Navarro et al., 2008; Payer et al., 2013). As a mirror image to the downregulation of *Xist,* we observed the upregulation of *Tsix* (**Figure 5B**), the antisense repressor of *Xist* during X-inactivation (Lee and Lu, 1999; Luikenhuis et al., 2001; Sado et al., 2001). However, as we have been using a functionally null *Tsix* truncation (TST) allele on the X_mus_ in our study (Luikenhuis et al., 2001; Ogawa et al., 2008), we confirmed previous findings that *Tsix* is dispensable for *Xist* downregulation during X-reactivation in iPSCs (Maclary et al., 2014; Payer et al., 2013).

Next, we investigated the regulatory landscape of the *Xic* from a structural perspective. The *Xic* is divided into two functionally opposing domains (van Bemmel et al., 2019; Galupa et al., 2020; Nora et al., 2012; Spencer et al., 2011; Tsai et al., 2008): TAD-D, which contains the non-coding genes *Tsix, Xite* and *Linx,* which are repressors of *Xist*; and TAD-E, which harbors *Xist* itself and its activators, the non-coding *Jpx* and *Ftx* and the protein-coding *Rlim/Rnf12* (**Figure 5C**). First, we noticed a gradual strengthening of the TAD border between TAD -D and TAD-E during reprogramming, as indicated by a drop in the insulation score (**Figure 5C**). Then we focussed on the structural organization of TAD-D. Whereas changes in expression of *Tsix* and *Linx* already occurred early during reprogramming (**Figures 5B and 5D**), we could not detect major changes in the 3D-organisation of TAD-D on the inactive X_mus_ at D5 (**Figures 5C, 5E, 5F, and 5G**) despite seeing them on the active X_cas_ (**Figures S5C-E**). This shows that restructuring of TAD-D only occurs after X-reactivation, suggesting it to be a consequence rather than a cause of *Xist* downregulation.

On the contrary, when we assessed the organization of TAD-E, we noted various changes early on. We observed a decrease in 3D-interactions on the Xi (X_mus_) within the region spanning *Xist*, *Jpx,* and *Ftx* already on D5 preceding X-reactivation, while we saw an increase in interactions between *Ftx* and *Rlim* (**Figures 5C-5G**). The kinetics in expression changes of the Xist activators *Ftx* (Furlan et al., 2018) and *Rlim* (Gontan et al., 2012; Jonkers et al., 2009) did not provide a clear correlation with Xist downregulation, making them unlikely candidates for *Xist* downregulation in our system. However, *Jpx,* which interacts with *Xist in cis* (Tsai et al., 2008) and facilitates *Xist* expression during X-inactivation (Sun et al., 2013; Tian et al., 2010), was downregulated with a highly similar profile to *Xist* during reprogramming, however, slightly preceding it. This suggests that *Jpx* downregulation might play a facilitative role in decreasing *Xist* levels during X-reactivation.

We conclude that the *Xic* is remodeled during reprogramming at multiple levels, leading to downregulation of *Xist*, a critical step for X-reactivation. We observed early structural changes in *Xist* TAD-E, where we detected an early loss of regulatory contacts in between *Xist* and its activators *Jpx* and *Ftx*. Finally, downregulation of *Jpx* and changes in chromatin accessibility at the pluripotency factor bound *Xist* intron 1 and the *Xist* promoter preceded *Xist* downregulation, suggesting them as potential candidate mechanisms to be involved in the process.

### Structural Remodeling of the X Chromosome Occurs in the Absence of Chromatin Opening and Reactivation of Transcription

The relationship and interplay between chromosome architecture and transcription during development has been an area of intense debate (Lucic et al., 2019; Rowley and Corces, 2018). Whereas some evidence suggests that transcription determines 3D chromatin organization (Rowley et al., 2017), it has been shown previously, that formation of TADs during zygotic genome activation is independent of transcription, and not merely a consequence of it (Du et al., 2017; Hug et al., 2017; Ke et al., 2017). While there might be context-dependent differences, the inactive X illustrates a unique instance, where TADs are actively attenuated (Wang et al., 2018) by the action of the non-coding Xist RNA and are fully regained during X-reactivation. To get more mechanistic insights, we thus wanted to ask if the formation of structural domains on the inactive X precedes or rather follows chromatin opening and gene reactivation.

When inspecting Hi-C matrices on D5, we noticed a unique intermediate structure of the inactive X (**Figures 2A and 6A**). It showed typical Xi features such as the mega-domains and their associated super-loops (Darrow et al., 2016) between *Dxz4* and *Firre*, as well as between *x75* and *Dxz4* (**Figure S6A**), but also already emerging TAD structures typical of an active X (**Figure 6A**). To assess the gain of TADs during X-reactivation on a quantitative level, we computed the insulation score (Crane et al., 2015) to assess the strength of TAD borders, and the domain score (Krijger et al., 2016) to quantify the degree of connectivity within TADs. In contrast to compartments, we already noticed changes in both insulation score and domain score in D5 P-RFP+ cells (**Figures 6B, 6C, and 6D**). Specifically, we observed a strengthening of TAD borders (**Figure 6B**) and an increase in the range of the insulation score (**Figures 6C** **and S6B**). Moreover, we detected a significant increase in the domain score of the Xi (X_mus_) (**Figures 6D** **and S6C**), revealing an increased connectivity of TADs. Our observation of preferential binding of Xist RNA to gene-rich A-like subcompartments (**Figure 3F**) and its role in repelling architectural proteins like CTCF and cohesin from the inactive X (Minajigi et al., 2015), prompted us to ask if this would lead to differential domain score dynamics among subcompartments. Indeed, when we assessed the relative domain score, to highlight changes occurring at D5, we observed that B-like subcompartments underwent the largest increase in TAD connectivity, while A-like compartments lagged behind (**Figures 6E** **and S6D**). Moreover, when we assessed domain score differences and correlated these to local enrichment of Xist RNA in NPCs, we found a strong anti-correlation between the levels of Xist, and the relative domain score at D5 (**Figure 6F**). This suggests that high levels of Xist inhibit the early formation of TADs, in agreement with the delayed gain in TADs observed in generally Xist-rich A-like compartments when compared with Xist-poor B-like compartments. Furthermore, considering that levels of H3K27me3 highly correlate with levels of Xist RNA, we expectedly observed a similar anticorrelation between H3K27me3 levels and domain score dynamics (**Figure 6G**).

**Figure 6.**
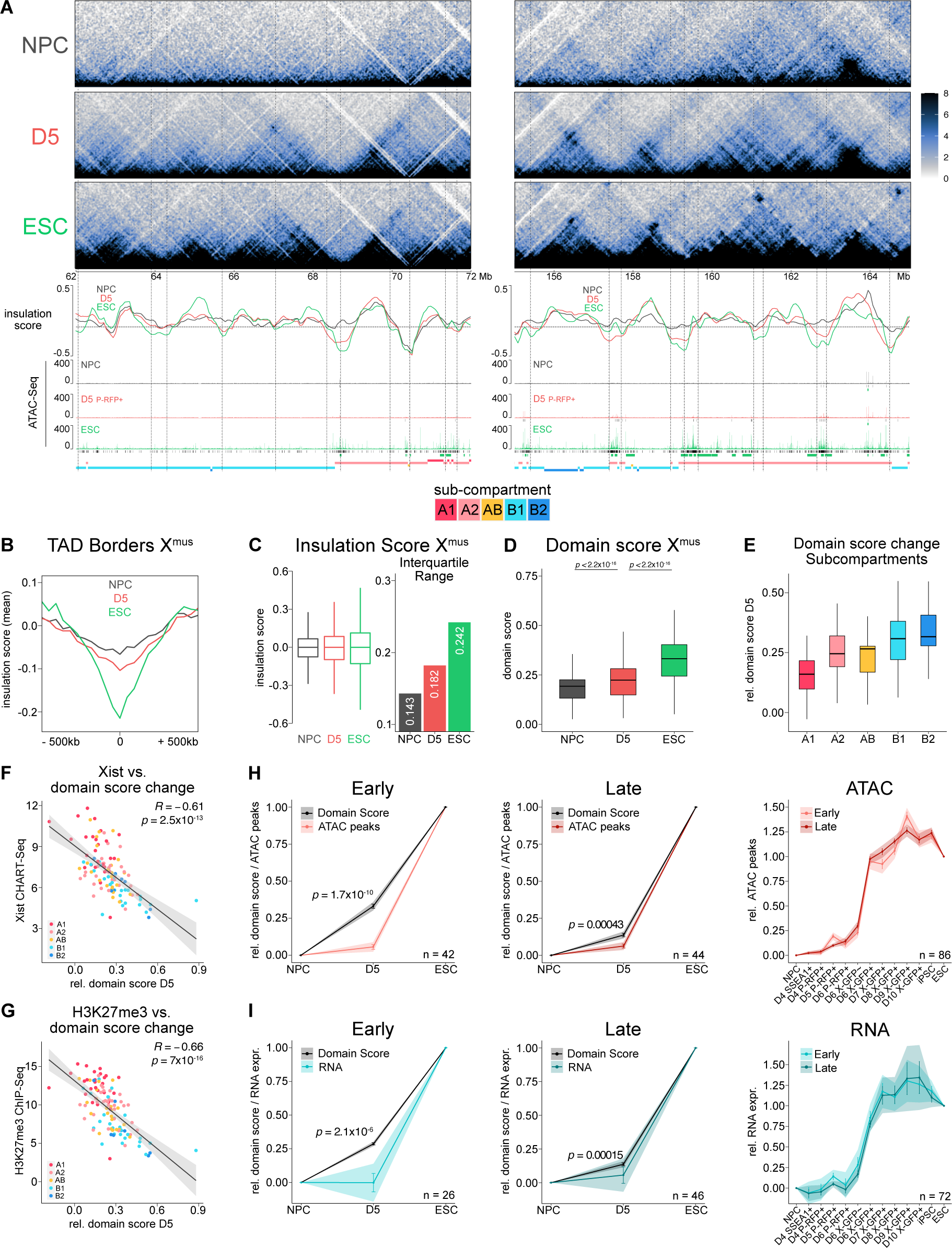
Structural Changes During X-Reactivation in the Absence of Chromatin Opening and Transcription. (A) Two representative X-linked regions of 10 Mb for early TAD formation are shown. Allele-specific Hi-C map of chromosome X_mus_ at 20-kb resolution. Scale is shown in mega-bases (Mb). Insulation scores at 50-kb resolution, dashed line indicates cut-off for TAD borders at -0.086. ATAC-Seq profiles with ATAC peaks shown in black (except for NPCs, differential peaks compared to NPCs are shown). Genes either escaping X-inactivation in NPCs or being reactivated based on RNA expression are shown in green. Position of subcompartments is shown at the bottom. (B) Meta region plot of insulation score at TAD boundaries at each time-point (n = 116). Lines show mean. (C) Comparison of insulation scores for chromosome X_mus_. Interquartile range of insulation scores is shown on the right. (D) Comparison of domain scores for chromosome X_mus_. The p values are calculated by Wilcoxon rank-sum test. (E) Degree of change in domain score of X_mus_ of subcompartments on D5. The relative domain score at D5 = (D5-NPC)/(ESC-NPC). (F) Correlation between Xist RNA CHART-Seq enrichment in NPCs (Wang et al., 2018) and the relative domain score at D5 is shown. Points represent TADs. Colors of points indicate subcompartments. R and p values calculated by Pearson’s correlation are shown. Black line represents linear regression fitting. Shading denotes 95% confidence interval of the fit. (G) As (F) for H3K27me3 ChIP-Seq in NPCs (Wang et al., 2018). (H) Comparison of domain dynamics and chromatin opening. Relative domain score is shown. Relative sum of ATAC peaks per TAD is shown. Only TADs with a minimum of 15 peaks in ESC were used. Early, TADs that changed from NPC to D5 (**Figure S6E**). Late, TADs that did not change from NPC to D5. Line shows mean. Error bars denote SEM. Shading denotes 95% confidence interval. The p values are calculated by Wilcoxon rank-sum test. Right panel shows comparison of chromatin opening dynamics (relative ATAC peaks) between early and late TADs. (I) Comparison of domain dynamics and gene reactivation. Relative domain score is shown. Relative mean expression per TAD is shown. Early, TADs that changed from NPC to D5. Late, TADs that did not change from NPC to D5. Line shows mean. Error bars denote SEM. Shading denotes 95% confidence interval. The p values are calculated by Wilcoxon rank-sum test. Right panel shows comparison of gene reactivation dynamics (relative RNA expression) between early and late TADs.

To identify TADs that have undergone domain score changes between stages, we performed *k*-means clustering on the relative domain score (**Figure S6E**), which showed that 55% of TADs already increased connectivity at D5 by more Whan 20% (“early”, increase in domain score; “late”, no increase in domain score). In line with our previous observations, we found B-like subcompartments to be enriched in early TADs, whereas TADs of subcompartment A1 were mostly designated late TADs (**Figure S6F**). When we then assessed chromatin opening at early TADs, we observed a significantly higher degree in domain score change, compared to an only mild increase in the number of ATAC peaks suggesting that TAD formation occurs before the appearance of chromatin opening (**Figures 6A and 6H**). In agreement with this and our observation that chromatin opening precedes gene reactivation (**Figures S4H and S4I**), we also found that TAD formation occurred in the absence of significant changes in gene expression at D5 (**Figure 6I**). Moreover, when we compared the kinetics of chromatin opening (**Figure 6H**) and gene reactivation (**Figure 6I**) between early and late TADs, we did not find any significant differences. Therefore we conclude that TAD formation does not necessa rily direct chromatin opening and gene reactivation.

Taken together, we show that early changes in TAD connectivity initiate from B-like subcompartments on the inactive X and anti-correlate with the local presence of Xist RNA. Moreover, our data show that TAD formation during X-reactivation often precedes and occurs without significant chromatin opening and gene reactivation, while intriguingly early TADs do not open chromatin or reactivate genes before late TADs. This suggests that chromatin opening and transcription are not essential drivers of the structural remodeling of the X chromosome during X-reactivation or vice versa, illustrating the mechanistic independence between these two events.

## Discussion

How changes in chromatin conformation and transcriptional activity are interlinked during cell fate transitions has been a topic of intense debate (Rowley and Corces, 2018; Stadhouders et al., 2019). Here we have used X-chromosome reactivation during iPSC-reprogramming as a model system to address this question, as it allowed us to study the chromosome -wide switch from an inactive, heterochromatic state into an active, euchromatic one. We thereby uncovered an underappreciated A/B-like compartment structure on the inactive mouse X chromosome, which resembles its active counterpart and separates distinct chromatin domains. We detect the first signs of X-reactivation to initiate from regions escaping XCI, while full reactivation of most genes occurred in a switch-like fashion, coinciding with the downregulation of Xist RNA. TAD structures emerged during X-reactivation before apparent gene reactivation, suggesting that transcriptional and structural remodeling of the X chromosome are independently suppressed by Xist RNA, and therefore qualify as functionally distinct events.

### Rapid Reactivation of the Inactive X-Chromosome

Previous studies on X-reactivation dynamics during iPSC reprogramming were based on mouse embryonic fibroblast (MEF) reprogramming systems (Janiszewski et al., 2019; Pasque et al., 2014; Payer et al., 2013). These suffered from low X-reactivation efficiencies and high sample heterogeneity, with only a small fraction of cells at a given time point being poised to undergo X-reactivation. This resulted in slow, gradual and asynchronous X-reactivation kinetics, lasting over the course of several days, making it difficult to study its steps on a regulatory level.

We therefore developed PaX, a tailor-made iPSC-reprogramming system based on a dual pluripotency and X-reporter mouse ESC line that allowed us to obtain for the first time a high-resolution time course of gene reactivation and chromatin opening during X-reactivation and intersected it with changes in 3D-chromatin structure. Our system allowed us to isolate large amounts of homogeneous cell populations poised for X-reactivation, which would subsequently progress synchronously with near-deterministic efficiency through X-reactivation, enabling us to faithfully analyze the stepwise progression of X-reactivation during reprogramming (**Figure 1**). Therefore, when analyzing allele-specific gene expression dynamics of the inactive X, we could demonstrate that X-reactivation in our system occurred rapidly in the time span of approximately 24 hours, mirroring faithfully the kinetics of the X-reactivation process *in vivo* in mouse blastocysts (Borensztein et al., 2017).

### Compartmentalization of the Inactive X-Chromosome

A prominent feature of eukaryotic chromosomes is their spatial segregation into two compartments: A, corresponding to open chromatin and high mRNA expression, and B, corresponding to closed chromatin and low expression (Lieberman-Aiden et al., 2009), which manifest themselves as a distinct checkerboard pattern on Hi-C matrices. However, the mouse inactive X-chromosome has served as a unique exception to this observation, as it has been described to be devoid of A/B compartments and to be organized instead into two large mega - domains (Giorgetti et al., 2016; Wang et al., 2018, 2019). Here, we have uncovered that the mouse inactive X-chromosome is in fact segregated into A/B-like compartments that resemble the A/B structure on the active X within NPCs (**Figure 2**). While the high resolution of our Hi-C samples has facilitated this discovery, we show that this has been previously overlooked due to the strong features of the two mega-domains. The mega-domains are predominant when applying principal component analysis on Hi-C correlation matrices of the Xi, thereby obscuring the underlying compartment structure, which could only be unveiled by our separate analysis of the two mega-domains. Similarly, PCA on human chromosomes 4 and 5 initially was onl y able to capture the p and q arms, and only after splitting at the centromere, was the underlying A/B compartment structure revealed (Lieberman-Aiden et al., 2009). Moreover, our findings are supported by observations made in human cells, where compartments on the inactive X chromosome have been observed previously (Darrow et al., 2016), and mouse primary neurons, where compartments have been suggested to exist on the Xi as well. Furthermore, when we applied our analysis strategy of performing PCA separately for each mega-domain to published NPC and MEF *in situ* Hi-C data (Wang et al., 2018, 2019), we also observed A/B-like compartment structures on the Xi (**Figure S2**). This suggests that the underlying A/B-like compartmentalization is a general biological feature of the inactive X chromosome, which might have been overlooked in previous studies for technical reasons. Importantly, we note that our observation of A/B-like compartments on a transcriptionally inactive chromosome favors a model where compartmentalization is not always driven by gene transcription.

It will therefore be of particular interest to identify the underlying principles shaping compartment structures, for example, if the mutually exclusive enrichment of repeated sequences like SINEs (in A-compartment) or LINEs (in B-compartment) may play an instructive role (Lu et al., 2019). One notable aspect of the A/B-like compartments on the Xi is their distinct chromatin status, with A-like compartments being enriched in Xist RNA and the Polycomb-based H3K27me3 mark when compared to B-like compartments showing higher levels of CBX1, a reader of H3K9 dimethylation (**Figure 3**). Nevertheless, except for escapees in the A-like compartment, both A-like and B-like compartments are transcriptionally inactive on the Xi. The Xi’s A-like compartments differ from the classical A compartments present on the active X, which are active in transcription and enriched in H3K4me3, despite showing structurally similar interaction patterns. This could be explained by the fact that Xist RNA and Polycomb proteins during X-inactivation first enter into strongly transcribed, gene-rich A-type regions (Pinter et al., 2012; Simon et al., 2013), while only later and less efficiently also spreading into gene-poor, already silent B-type heterochromatin (Pinter, 2016). This makes sense from a functional point of view, where Xist-based silencing would be predominantly needed in the actively transcribed A-compartment to achieve X-linked gene dosage compensation while being less critical for gene -poor H3K9me-marked heterochromatin in the B-compartment. As a consequence, Xist and its interacting partners might stabilize this structure, thereby establishing an epigenetic memory of the original A-compartment structure present at the time of X-inactivation, which is then maintained on the inactive X chromosome. The spatial separation between two types of heterochromatin on the inactive X, the Xist/Polycomb-rich A-like and the H3K9me-rich B-like heterochromatin, could be driven by liquid-liquid phase separation (LLPS) mechanisms, which have been previously described for both Polycomb (Plys et al., 2019) and H3K9me2/3 domains (Larson et al., 2017; Strom et al., 2017). Indeed, Xist RNA recruits a multitude of factors involved in LLPS to the inactive X (Cerase et al., 2019) and Xist-deletion or depletion of Xist-associated LLPS factors like PRC1 and hnRNPK has been shown to significantly compromise the underlying Xi compartment structure (Wang et al., 2019). Although A/B-like compartments of the inactive X, similar to the previously described S1 (A-like) and S2 (B-like) compartments (Wang et al., 2018, 2019), were thought to only exist transiently during X-inactivation, before subsequently being merged by SMCHD1, we were able to show that the compartmentalization persists on the inactive X and can be unveiled when applying separate analysis to the two-mega-domains. In support of our data, this dual heterochromatin structure has been previously observed on the human Xi (Chadwick and Willard, 2004), where it coincides with compartment structures as well (Darrow et al., 2016), suggesting it to be a conserved property of the inactive X chromosome in mammals.

### Spatial Sub-Megabase Clusters and their Role in Early X-Reactivation

Underlying an A/B-like compartment structure, we have additionally discovered distinct spatial clusters and subcompartments on the sub-megabase level (**Figure 3**). Intriguingly, we found that gene-rich clusters, among others, are characterized by close spatial proximity and preferential binding of Xist RNA, arguing that spatial clustering of the inactive X provides a 3D-scaffold for efficient Xist-mediated gene silencing and dosage compensation. This is in line with observations that these gene-rich domains are the first areas to be coated by Xist RNA during the X-inactivation process (Simon et al., 2013).

Furthermore, we noticed that while the timing of complete X-linked gene reactivation was conserved in all subcompartments, partial X-reactivation early in reprogramming could be observed in a distinct spatial cluster, characterized by a high density in genes escaping X - inactivation (**Figure 4**), complementary to a previous observation that regions of high escapee density coincide with TADs in ESCs (Marks et al., 2015). This suggested that proximity to accessible chromatin at escapees might facilitate further opening of regions nearby in a zipper-like fashion, leading to a partial basal reactivation of these genes. Indeed, we found this specific subset of early partially reactivated genes to lie in close proximity to escapees. However, the timing of complete X-reactivation of these genes was conserved compared to the rest of the genes. This suggests that while close distance to escapees, as reported previously (Janiszewski et al., 2019), plays a role in X-reactivation, different mechanisms regulate the timing of complete X-reactivation, as discussed below.

### *Xist* Downregulation: The Key Step for X-Reactivation Timing

An important event both for X-linked gene reactivation as well as for the structural remodeling of the X chromosome is the downregulation of *Xist* expression. Xist RNA has multiple distinct roles in establishing the silent chromatin state during X-inactivation based on its interaction with critical architectural and silencing factors like Cohesin, or SHARP/SPEN (Chu et al., 2015; McHugh et al., 2015; Minajigi et al., 2015). Therefore this is in line with our observation that high Xist RNA occupancy on the inactive X anticorrelated with early TAD formation (**Figures 6 and S6**) and that gene reactivation kinetics were tightly linked to a sharp drop in *Xist* levels (**Figures 1 and 5**). It is well appreciated that *Xist* is both, directly and indirectly, repressed by pluripotency factors such as OCT4, SOX2, NANOG, PRDM14, and ZFP42/REX1 (Payer and Lee, 2014). Indeed, we found that *Xist Intron 1*, a known pluripotency factor binding hub (Navarro et al., 2008; Payer et al., 2013), rapidly gained in accessibility with expression of the MKOS reprogramming cassette, coinciding with a partial decrease in *Xist* promoter accessibility and in Xist RNA levels early on during reprogramming (**Figures 1 and 5**). This might allow the initial partial reactivation of genes lying near escapees, which we observed in structural cluster 5 (**Figures 4 and S4**). However, with the expression of endogenous naive pluripotency factors on D6 of reprogramming (**Figure 1**), we saw a drastic drop in *Xist* expression, in line with NANOG, ZFP42 and PRDM14 being important *Xist* repressors (Gontan et al., 2012; Navarro et al., 2008; Payer et al., 2013). As this coincided temporally with the full reactivation of X-linked genes, our data suggest that *Xist* downregulation is indeed the rate-limiting step during X-reactivation thereby coupling it functionally to the reprogramming process.

Apart from *trans*-regulation by pluripotency factors, the *Xist* locus is regulated locally at the X-inactivation center by the activators *Jpx*, *Ftx,* and *Rlim* within *Xist* TAD-E and repressors like *Tsix*, *Xite* and *Linx* within its neighboring TAD-D (van Bemmel et al., 2019). When we assessed the topology of both TADs during reprogramming (**Figures 5 and S5**), we observed early structural changes to occur in particular within *Xist* TAD-E, suggesting it to be a main driver in *Xist* downregulation. Especially suggestive was the downregulation of *Jpx*, which slightly preceded the kinetics of *Xist*-downregulation. *Jpx*, being a critical activator of *Xist* during X-inactivation (Sun et al., 2013; Tian et al., 2010), might therefore also play a role in *Xist* downregulation during reactivation. How *Jpx* itself is regulated and what might be its functional impact on *Xist* during X-reactivation remain open questions warranting further analysis.

### Early TAD Formation Occurs in the Absence of Chromatin Opening and Reactivation of Transcription

Here we have shown that during X-reactivation, chromatin opening and gene reactivation are tightly linked (**Figures 4 and S4**). However, it was unknown whether changes in chromatin conformation are closely connected, as previously shown for autosomes during iPSC reprogramming (Stadhouders et al., 2018). Moreover, we demonstrate that as the human Xi, the mouse inactive X chromosome features an A/B-like compartment structure (**Figure 2**), and consistent with previous findings (Wang et al., 2018) we found TADs to be strongly attenuated on the Xi (**Figure 6**). This highlights that compartmentalization and TAD organization depend on distinct mechanisms (Nora et al., 2017; Schwarzer et al., 2017) and that TADs need to be fully reestablished during X-reactivation. Intriguingly, a recent study on the dynamics of TADs during imprinted X-inactivation suggested that loss of TAD structure rather follows gene silencing and further showed that maintenance of TADs was restricted to escapee regions (Collombet et al., 2020), which would propose a model where dynamics of TADs and transcription are intertwined on the Xi.

When we analyzed the Xi domain structure at day 5 of reprogramming, one day before the onset of full transcriptional reactivation, we found that more than 50% of TADs already displayed a significant increase in TAD connectivity (**Figures 6 and S6**). Considering our observation that mega-domains are still present at D5 (**Figure 2**), this shows that TADs and mega-domains are distinct structural entities that are controlled independently. This is in line with the disappearance of TADs in the absence of mega-domain formation during imprinted X-inactivation (Collombet et al., 2020) and the observation that mega-domains are dispensable for gene silencing and TAD attenuation during X-inactivation (Bonora et al., 2018; Froberg et al., 2018; Giorgetti et al., 2016).

Moreover, when we then compared these changes to the dynamics of chromatin accessibility and gene reactivation, we found that the connectivity of early TADs had already undergone significantly higher changes, compared to relatively mild changes in both chromatin accessibility and gene reactivation. This suggests that chromatin opening and transcription are not essential drivers of the structural remodeling of the Xi. Moreover, early TAD formation did not seem to prime for early reactivation either, as early and late TADs shared highly similar gene reactivation and chromatin opening dynamics, illustrating that these are mechanistically separate events, and that TADs are not required for transcription as shown previously (Nora et al., 2017). However, a common denominator of both processes seems to be Xist. When we assessed the enrichment of Xist RNA, we noticed a strong correlation of early TAD formation with low Xist RNA occupancy, therefore early TADs forming mostly in Xist-poor B-like compartments and late TADs in Xist-rich A-like compartments. Considering Xist’s role in repelling architectural proteins like CTCF and cohesin from the inactive X (Chu et al., 2015; Minajigi et al., 2015), we propose that low levels of Xist facilitate the early restructuring of TADs, by allowing increased binding of CTCF and cohesin. Xist’s gene silencing function however depends on a different set of binding partners than its structural role (Chu et al., 2015; McHugh et al., 2015; Minajigi et al., 2015). The removal of Xist-dependent repressive chromatin marks like H3K27me3 by UTX/KDM6A (Borensztein et al. 2017) and the gain in histone acetylation on the Xi after Xist downregulation (Janiszewski et al., 2019), in combination with Xist-independent DNA-demethylation (Pasque et al., 2014), have been shown to facilitate the transcriptional reactivation of the X. Therefore it is not surprising that this multistep process does not follow the same kinetics as the gain of TADs, although both processes are controlled by Xist, again highlighting its key role during X-reactivation.

### Conclusions & Outlook

Overall, our study provides mechanistic insight into the process of X-reactivation and the long-debated relationship of genome topology and transcription. We provide evidence that the mouse inactive X chromosome is in fact not as “unstructured” as it was believed to be and that, together with Xist downregulation, the fine structure of the inactive X parallels the reactivation kinetics. Moreover, our comprehensive dataset of the dynamics of transcriptional reactivation and chromatin opening and our high-resolution chromosome conformation maps of the reactivating X will provide a useful resource for future studies. Finally, our tailor -made PaX reprogramming system constitutes an optimized framework for further analysis of the chromatin dynamics and functional dissection of the X-reactivation process and, more generally, of the interplay between chromosome organization, chromatin architecture and gene regulation.

## Acknowledgments

We thank J. Lee for EL16.7 TST ES cells; K. Kaji for reprogramming cassette plasmids; J. Nathans for X-GFP plasmids; R. Stadhouders and G. Stik for advice on Hi-C technology; P. Soler for advice on bioinformatic analyses; the CRG Genomics Unit for sequencing; the CRG/UPF FACS Unit for FACS sorting; and members of B.P.’s laboratory for discussions. We also thank R. Stadhouders and J. Valcarcel for critical reading of the manuscript. This work was supported by the European Research Council under the 7th Framework Programme FP7/2007-2013 (ERC Synergy Grant 4D-Genome, grant agreement 609989 to G.J.F.), by the Spanish Ministry of Science, Innovation and Universities (BFU2014-55275-P, BFU2017-88407-P to B.P. and PGC2018-099807-B-I00 to G.J.F.), the Agencia Estatal de Investigación (AEI) (EUR2019-103817 to B.P.), the AXA Research Find (to B.P.) and the Agencia de Gestio d’Ajuts Universitaris i de Recerca (AGAUR, 2017 SGR 346 to B.P.) and by the NIH grant R35GM123926 to S.F.P.. We would like to thank the Spanish Ministry of Economy, Industry and Competitiveness (MEIC) to the EMBL parWnerVhip and to the “Centro de Excelencia Severo Ochoa”. We also acknowledge support of the CERCA Programme of the Generalitat de Catalunya. M.B. was supported by a La Caixa International PhD Fellowship.

## Author Contributions

M.B. and B.P. conceived the study and wrote the manuscript with input from all coauthors; M.B. established reporter cell line, performed reprogramming experiments and performed molecular biology, RNA-Seq, ATAC-Seq, and *in situ* Hi-C experiments; M.B., E.V., E.Z., and G.J.F. performed bioinformatic analyses with input from S.F.P.; M.B. and E.V. integrated and visualized data; B.P. and G.J.F. acquired funding and supervised the research.

## Experimental Model and Subject Details

### Embryonic Stem Cell Culture

Mouse embryonic stem cells (ESCs) were cultured on 0.2% gelatin-coated dishes in DMEM (Thermo Fisher Scientific, 31966021), supplemented with 10% FBS (ES-qualified, Thermo Fisher Scientific, 16141079) 1,000 U/ml LIF (ORF Genetics, 01-A1140-0100), 1 mM Sodium Pyruvate (Thermo Fisher Scientific, 11360070), 1x MEM Non-Essential Amino Acids Solution (Thermo Fisher Scientific, 11140050), 50U/ml penicillin/streptomycin (Ibian Tech, P06-07100) and 0.1 mM 2-mercaptoethanol (Thermo Fisher Scientific, 31350010). Cells were incubated at 37°C with 5% CO2. The medium was changed every day and cells were passaged using 0.05% Trypsin -EDTA (Thermo Fisher Scientific, 25300054). Cells were monthly tested for mycoplasma contamination using PCR.

## Generation of Cell Lines

### X-GFP Reporter

We used the female F2 ESC line EL16.7 TST, that was derived from a cross of *Mus musculus musculus* with *Mus musculus castaneus* (Ogawa et al., 2008). As a result, cells contain one X chromosome from *M.m musculus* (X_mus_) and one from *M.m castaneus* (X_cas_). Moreover, EL16.7 TST contains a truncation of *Tsix* on X_mus_ (*Tsix*TST/+), which abrogates *Tsix* expression and leads to the non-random inactivation of X_mus_ upon differentiation.

A GFP reporter construct was targeted into the second exon of *Hprt* on X_mus_ as follows: Homology arms flanking the target site were amplified from genomic DNA and cloned into pBluescript II SK(+) (Addgene, 212205) by restriction-enzyme based cloning and the cHS4-CAG-nlsGFP-cHS4 construct, kindly provided by J. Nathans (Wu et al., 2014), was cloned between the two homology arms.

5×10^6^ cells were mixed with 1.6 μg circularised targeting vector and 5 μg single guide RNA vector PX459 (Addgene, 48139) (5’-TATACCTAATCATTATGCCG-3’), to achieve an optimal ratio of Cas9 to targeting vector equal to 5:1 (Pinder et al., 2015). Cells were nucleofected with the AMAXA Mouse Embryonic Stem Cell Nucleofector Kit (LONZA, VPH-1001) using program A-30 and 7.5 μM RS-1 (Merck, 553510) was added to enhance homology-directed repair. To select for the disruption of *Hprt*, cells were grown in the presence of 10 μM 6-thioguanine (Sigma-Aldrich, A4882-250MG) for 6 days, and GFP+ cells were isolated by FACS using a BD Influx (BD Biosciences). Single clones were screened by Southern blot hybridization. Inactivation of the X - GFP construct upon differentiation was confirmed using embryoid body differentiation.

pSpCas9(BB)-2A-Puro (PX459) was a gift from Feng Zhang (Addgene plasmid # 48139; http://n2t.net/addgene:48139; RRID:Addgene_48139)

### Reprogramming Cassette

An all-in-one gene targeting vector with doxycycline-inducible reprogramming factors, MKOSimO neotk rtTA Sp3, kindly provided by K. Kaji (Chantzoura et al., 2015), was targeted into the third intron of the *Sp3* gene in the ESC line EL16.7 TST X-GFP using CRISPR-Cas9. 5×10^6^ cells were mixed with 3.8 μg circularised targeting vector and 2.5 μg single guide RNA vector PX459 (5’-GTGACAATCTCCGGAAAGCG-3’) and nucleofected with the AMAXA Mouse Embryonic Stem Cell Nucleofector Kit (LONZA, VPH-1001) using program A-24. 7.5 μM RS-1 (Merck, 553510) was added to enhance homology-direcWed repair. Cells were selected with 300 μg/ml G418 for 5 days. Clones were selected for expression of mOrange upon the addition of 1 mg/ml doxycycline for 24 hours and then screened by Southern blot hybridization.

Knockout of mOrange was generated using CRISPR-Cas9. 5×10^6^ cells were mixed with 1.8 μg single guide RNA vector PX459 V2 (Addgene, 62988) (5’-CAACGAGGACTACACCATCG-3’) and nucleofected with the AMAXA Mouse Embryonic Stem Cell Nucleofector Kit (LONZA, VPH -1001) using program A-30. mOrange-negative cells were isolated by FACS using a BD Influx and single clones were screened for maintenance of proper cassette expression by quantitative RT - PCR.

pSpCas9(BB)-2A-Puro (PX459) V2.0 was a gift from Feng Zhang (Addgene plasmid # 62988; http://n2t.net/addgene:62988; RRID:Addgene_62988)

### Southern blot

Genomic DNA (10 μg) was digested with appropriate restriction enzymes overnight. Subsequently, genomic DNA was separated on a 0.8% agarose gel and transferred to an Amersham Hybond-XL membrane (GE Healthcare, RPN303S). Probes were synthesized by PCR amplification and labeled with dCTP, [α-32P] (Perkin Elmer, NEG513H250UC) using High Prime (Roche, 11585592001) and hybridization performed in Church buffer.

### P-RFP Pluripotency Reporter

Lentivirus, encoding the mouse *Nanog* promoter driving RFP expression, was purchased from System Biosciences (SR10044VA-1). EL16.7 TST X-GFP MKOS ESCs were infected at an MOI of 30 and RFP-positive cells FACS purified using a BD Influx. Single clones were isolated and selected based on proper RFP expression using FACS analysis on a BD LSRFortessa.

### Neural Precursor Cell Differentiation

ESCs were differentiated to neural precursor cells (NPCs) as described previously (Abranches et al., 2009). ESCs were seeded at a density of 2.75×10^5^ cells/cm^2^ in N2B27 (50% DMEM/F12 (Thermo Fisher Scientific, 21041025), 50% Neurobasal medium (Thermo Fisher Scientific, 12348017), 1x N2 (Thermo Fisher Scientific, 17502048), 1x B27 (Thermo Fisher Scientific, 12587001)) supplemented with 0.4 μM PD0325901 (Selleck Chemicals, S1036-5mg), 3 μM CHIR99021 (Sigma-Aldrich, SML1046-5MG) and 1,000 U/ml LIF (ORF Genetics, 01-A1140-0100). 24 hours later, cells were dissociated using Accutase (Thermo Fisher Scientific, 00-4555-56) and plated at 2.95×10^4^ cells/cm^2^ in RHB-A (Takara Bio, Y40001) on 0.2% gelatin-coated T75 flasks, changing media every other day. On days 6 and 8, media was supplemented with 10 ng/ml EGF (R&D Systems, 236-EG-200) and 10 ng/ml bFGF (Thermo Fisher Scientific, 13256029) and additionally with 10μM ROCK inhibitor (Sellekchem, S1049) on day 8. On day 9, cells were dissociated using Accutase (Thermo Fisher Scientific, 00-4555-56) and SSEA1 expressing cells were removed by MACS sorting using Anti-SSEA-1 (CD15) MicroBeads (Miltenyi Biotech, 130-094-530). To completely remove cells that hadn’t undergone XCI, cells were stained with SSEA-1 eFluor 660 (Thermo Fisher Scientific, 50-8813-42) for 15 min at 4°C, washed once with 0.5% BSA in PBS and then SSEA1-/Nanog-RFP-/X-GFP-cells were FACS purified using a BD FACSAria II SORP or a BD Influx (BD Biosciences) at a maximum flow rate of 4,000 ev/s to improve survival. FACS sorted cells were seeded at 3.5×10^5^ cells/cm^2^ on 0.2% gelatin-coated dishes in RHB-A, supplemented with EGF, FGF, and ROCKi. The medium was changed daily until day 12 when cells reached 100% confluency.

### Reprogramming of Neural Precursor Cells

Reprogramming of day 12 neural precursor cells was induced by the addition of 1 mg/ml doxycycline (Tocris, 4090/50) and 25 mg/ml L-ascorbic acid (Sigma-Aldrich, A7506-25G) to the NPC medium (RHB-A supplemented with EGF and FGF). 24 hours later, cells were dissociated using Accutase and seeded on irradiated mouse embryonic fibroblasts (iMEF) in ESC medium containing 15% FBS and supplemented with 1 mg/ml doxycycline and 25 mg/ml L-ascorbic acid. The medium was changed every other day. To isolate iPSC, SSEA1+/Nanog-RFP+/X-GFP+ cells were isolated using FACS at day 10 of reprogramming and re-plated on 0.2% gelatin-coated plates in ESC medium and kept in doxycycline free conditions for 5-6 days.

### RNA Isolation, Quantitative RT-PCR and RNA-Sequencing

RNA was extracted using the RNeasy Plus Mini Kit (Qiagen, 74136) or RNeasy Micro Kit (Qiagen, 74004) and quantified by Nanodrop. cDNA was produced with a High-Capacity RNA-to-cDNA Kit (Thermo Fisher Scientific, 4387406) and was used for qRT-PCR analysis in triplicate reactions with Power SYBR Green PCR Master Mix (Thermo Fisher Scientific, 4367659). Libraries were prepared using the TruSeq Stranded Total RNA Library Preparation Kit (Illumina, 20020597) followed by paired-end sequencing (2 x 125 bp) on an Illumina HiSeq 2500.

### Assay for Transposase-Accessible Chromatin with High Throughput Sequencing (ATAC-Seq)

ATAC-Seq was performed as described previously (Corces et al., 2017) with minor modifications. 50,000 FACS pXrified cellV Zere reVXVpended in 50 μl cold lysis buffer (10 mM TriV -HCl pH 7.4, 10 mM NaCl, 3 mM MgCl_2_, 0.01% Digitonin, 0.1% Tween-20, 0.1% IGEPAL CA-630). After 3 min the lysis was washed out using 1 ml cold lysis buffer containing Tween -20, but no Digitonin or IGEPAL CA-630. Cells were centrifuged for 10 min at 500 rcf and 4°C, supernatant was removed and nuclei were resuspended in 50 μl transposition reaction mix (25 μl Tn5 Transposase buffer (Illumina, 15027866), 2.5 μl Tn5 transposase (Illumina, 15027865), 16.5 —l PBS, 0.01% Digitonin, 0.1% Tween-20, 5 μl nuclease-free water) and incubated at 37°C for 45 min with 1000 RPM mixing. DNA was isolated using the MinElute PCR Purification Kit (Qiagen, 28004). Library amplification was performed by two sequential PCR reactions (8 and 4 -7 cycles, respectively) using the NEBNext High Fidelity PCR Master Mix (New England Biolabs, M0541S). DNA was then double-size selected using 0.5x and 1.5x Agencourt AMPure XP beads (Beckman, A63880) to isolate fragments between 100 bp and 1 kb. Library quality was assessed on a Bioanalyzer, followed by paired-end sequencing (2 x 125 bp) on an Illumina HiSeq 2500.

### Cell Isolation and Purification for ATAC-Seq and RNA-Seq

Cells were dissociated using Accutase (Thermo Fisher Scientific, 00-4555-56) (for NPCs), 0.05% Trypsin-EDTA (Thermo Fisher Scientific, 25300054) (for ESCs) or 0.25% Trypsin-EDTA (Thermo Fisher Scientific, 25200056) (for iPSCs) and then stained with SSEA-1 eFluor 660 (Thermo Fisher Scientific, 50-8813-42) for 15 min at 4°C. Cells were washed once with 0.5% BSA in PBS and then FACS sorted using a BD FACSAria II SORP or a BD Influx.

### Cell Isolation and Purification for Hi-C

Purification of G_0_G_1_ cells based on DNA content was performed as described previously (Bonev et al., 2017) with minor modifications. Briefly, cells were dissociated using Accutase (Thermo Fisher Scientific, 00-4555-56) (for NPCs), 0.05% Trypsin-EDTA (Thermo Fisher Scientific, 25300054) (for ESCs) or 0.25% Trypsin-EDTA (Thermo Fisher Scientific, 25200056) (for iPSCs) and then stained with SSEA-1 eFluor 660 (Thermo Fisher Scientific, 50-8813-42). iPSCs were additionally MACS sorted using Anti-SSEA-1 (CD15) MicroBeads (Miltenyi Biotech, 130-094-530). Cells were then fixed for 10 min at room temperature with freshly prepared 1% formaldehyde in PBS (SigA critical event during ma-Aldrich, F8775-4X25ML) and the reaction then quenched by addition of 0.2M glycine (NZYTech, MB01401). 1x10^6^ cells/ml were permeabilized using 0.1% saponin (Sigma-Aldrich, 47036-50G-F). 10 μg/ml DAPI (Thermo Fisher Scientific, D1306) and 100 μg/ml RNase A (Thermo Fisher Scientific, EN0531) were added and samples incubated for 30 min at room temperature protected from light with slight agitation. After washing once with cold PBS, samples were resuspended in cold 0.5% BSA in PBS at a concentration of 1x10^7^ cells/ml and immediately FACS purified using a BD FACSAria II SORP or a BD Influx. After FACS sorting, dry cell pellets were snap-frozen in dry ice and stored at -80°C.

### *In situ* Hi-C Library Preparation

*In situ* Hi-C was performed as described previously (Stadhouders et al. 2018) with minor modifications. One million cells purified for G_0_G_1_ were used as starting materials. Cells were lysed XVing 250 μl cold lysis buffer (10 mM TriV-HCl pH 8, 10 mM NaCl, 0.2% IGEPAL CA-630) supplemented with 50 μl protease inhibitor cocktail (Sigma-Aldrich, P8340-1ML). Cells were digested with 100 U MboI (New England Biolabs) and incubated for 2 hours at 37°C under rotation, followed by the addition of another 100U for 2 hours and another 100U before overnight incubation. The next day a final 100U were added and incubated for 3 hours. After fill-in with biotin-14-dATP (Thermo Fisher Scientific, 19524016), ligation was performed with 10,000 U T4 DNA Ligase (New England Biolabs, M0202M) overnight at 24°C under rotation. After de -crosslinking, DNA was purified using ethanol precipitation and sonicated to an average size of 300-700 bp with a Bioruptor Pico (Diagenode; seven cycles of 20 s on and 60 s off). Ligation products containing biotin-14-dATP were pulled-down using Dynabeads MyOne Streptavidin T1 beads (Thermo Fisher Scientific, 65601) and end-repaired and A-tailed using the NEBNext End Repair/dA-Tailing Module (New England Biolabs, E6060S and E6053S). Libraries were amplified using the NEBNext High Fidelity PCR Master Mix and NEBNext Multiplex Oligos for Illumina (New England Biolabs, M0541S and E7335S) for 8 cycles and size-selected with 0.9x Agencourt AMPure XP beads. Library quality was assessed on a Bioanalyzer and by low-coverage sequencing on an Illumina NextSeq 500, followed by high-coverage paired-end sequencing (2 x 125 bp) on an Illumina HiSeq 2500.

### RNA-Fluorescent *In Situ* Hybridization

Strand-specific RNA FISH was performed with fluorescently labeled oligonucleotides (IDT) as described previously (Del Rosario et al. 2017). Briefly, cells were fixed with 4% paraformaldehyde for 10 minutes at room temperature and then permeabilized for 5 minutes on ice in 0.5% Triton - X. 10ng/ml equimolar amounts of Cy5 labeled Xist probes BD384-Xist-Cy5-3’-AM (5’-ATG ACT CTG GAA GTC AGT ATG GAG /3Cy5Sp/ -3’) and BD417-5’Cy5-Xist-Cy5-3’-AM (5’-/5Cy5/ATG GGC ACT GCA TTT TAG CAA TA /3Cy5Sp/ -3’) were hybridized in 40% formamide, 10% dextran sulfate, 2xSSC pH 7 at room temperature overnight. Slides were then washed in 30% formamide 2xSSC pH 7 at room temperature, followed by washes in 2xSSC pH 7 and then mounted with Vectashield (Vector Laboratories, H1200). Images were acquired using an EVOS and a Cy5 light cube (Thermo Fisher Scientific).

## Quantification and Statistical Analysis

### Allele-Specific Analysis

Reads from the PaX hybrid cell line were disambiguated and mapped in two steps. First, each read was mapped in the reference genome of 129S1/SvImJ, and independently in the reference genome of CAST/EiJ (both genomes are available from the Wellcome Trust Sanger Institute ftp://ftp-mouse.sanger.ac.uk/REL-1504-Assembly) using BWA-MEM version 0.7.17 (with options -L 500 for ATAC-Seq and -t4 -P -k17 -U0 -L0,0 -T25 for Hi-C). In each case, the resulting SAM files were processed by samtools version 1.8 with the fixmate option to fill in mate coordinates. Each read was then assigned to a single genome and the coordinates were lifted over to the reference mm10 (*Mus musculus* genome of strain C57BL/6J) with custom Python scripts available for download at http://github.com/gui11aume/asmap. Briefly, genome assignment was carried out as follows: if the mapping quality of the read was 0 in each genome, the read was considered unmapped and no assignment was performed; otherwise, the read was assigned to the genome with the best alignment score (field AS:i from the output of BWA-MEM). When both alignment scores were equal, the genome of origin was considered ambiguous and no assignment was performed (but the coordinates of the read were still lifted over to mm10). Lift over to mm10 was also performed with custom Python scripts using the positions of the indels of 129S1/SvImJ and CAST/EiJ relative to C57BL/6J (Pinter et al., 2012). The positions of the indels relative to the reference strain C57BL/6J (mm10) were downloaded from ftp://ftp-mouse.sanger.ac.uk/REL-1505-SNPs_Indels/strain_specific_vcfs, and processed as explained in (Pinter et al., 2012).

### RNA-Seq Analysis

Mapping and disambiguation were performed as explained in the section Allele -Specific Analysis. Reads were mapped with STAR (Dobin et al., 2013) (standard options) and the Ensembl mouse genome annotation (GRCm38.78). Gene expression was quantified with STAR (--quantMode GeneCounts). Batch effects were removed using the ComBat function from the sva R package (v.3.22). Sample scaling and statistical analysis were performed with the R package DESeq2 (Love et al., 2014) (R v.3.3.2 and Bioconductor v.3.0), and the log_2_(vsd) (variance stabilized DESeq2) counts were used for further analysis unless stated otherwise. Standard TPM (transcripts per million) values were used as an absolute measure of gene expression.

### ATAC-Seq Analysis

Mapping and disambiguation were performed as explained in the section Allele -Specific Analysis. Peak calling was performed using Zerone (Cuscó and Filion, 2016), with option “make atac” at compile time. The ATAC-seq profiles were discretized using default parameters against the baseline set by the NPC profile. In the process, the four replicates of each time point were merged into a single discretized profile of resolution 300 bp. The NPC profile was discretized separately, with option “--no-mock” to indicate the absence of a separate baseline profile. In all cases, only the windows with a confidence score above 99.9% were considered to be ATAC-Seq peaks.

### *In situ* Hi-C data processing and normalization

Mapping and disambiguation were performed as explained in the section Allele -Specific Analysis, and only the read pairs where at least one end was mapped unequivocally were kept for further analyses (*i.e.*, with mapping quality greater than 0 and with a non-ambiguous chromosome of origin).

We processed Hi-C data by using an in-house pipeline based on TADbit (Serra et al., 2017). First, the quality of the reads was checked with FastQC to discard problematic samples and detect systematic artifacts. Trimmomatic (Bolger et al., 2014) with the recommended parameters for paired-end reads was used to remove adapter sequences and poor-quality reads (ILLUMINACLIP:TruSeq3-PE.fa:2:30:12:1:true; LEADING:3; TRAILING:3; MAXINFO:targetLength:0.999; and MINLEN:36). For mapping, a fragment-based strategy, as implemented in TADbit, was used, which is similar to previously published protocols (Ay et al., 2015). Briefly, each side of the sequenced read was mapped in full length to the reference genome (mm10, Dec 2011 GRCm38). After this step, if a read was not uniquely mapped, we assumed that the read was chimeric, owing to ligation of several DNA fragments. We next searched for ligation sites, discarding those reads in which no ligation site was found. The remaining reads were split as often as ligation sites were found. Individual split read fragments were then mapped independently. These steps were repeated for each read in the input FASTQ files. Multiple fragments from a single uniquely mapped read resulted in a number of contacts identical to the number of possible pairs between the fragments. For example, if a single read was mapped through three fragments, a total of three contacts (all-versus-all) was represented in the final contact matrix. We used the TADbit filtering module to remove noninformative contacts and to create contact matrices. The different categories of filtered reads applied were:

- Self-circle: reads coming from a single restriction enzyme (RE) fragment and point to the outside.
- Dangling-end: reads coming from a single RE fragment and point to the inside.
- Error: reads coming from a single RE fragment and point in the same direction
- Extra dangling-end: reads coming from different RE fragments but are close enough and point to the inside. The distance threshold used was left to 500 bp (default), which is between percentile 95 and 99 of average fragment lengths.
- Duplicated: the combination of the start positions and directions of the reads was repeated, pointing at a PCR artifact. This filter only removed extra copies of the original pair.
- Random breaks: start position of one of the reads was too far from the RE cutting site, possibly due to non canonical enzymatic activity or random physical breaks. Threshold was set to 750 bp (default), greater than percentile 99.9. From the resulting contact matrices, low-quality bins (those presenting low contacts numbers) were removed as implemented in TADbit’s ‘filter_columns’ routine. The matrices obtained were normalized for sequencing depth and genomic biases using OneD (Vidal et al., 2018). Then, they were further normalized for local coverage within the region (expressed as normalized counts per thousand within the region) without any correction for the diagonal decay. For differential analysis, the resulting normalized matrices were directly subtracted from each other.

To merge contact matrices, the counts of each replicate were summed in every 50 kb bin. Low mappability regions were excluded by removing rows / columns with read counts lower than 20% of the median. Finally, matrices were ICE-normalized using the implementation from the OneD R package (Vidal et al., 2018).

### Allele-Specific Expression

From a list of 806 protein-coding genes on the X-chromosome, we masked genes with insufficient sequence polymorphisms, leaving 558 genes. Moreover, to remove lowly expressed genes, we removed genes where expression from cas was below the 25th percentile, leaving 335 genes that passed these criteria. The allelic expression ratio of these genes was then calculated by dividing mus reads by the sum of mus and cas reads ((mus/mus+cas)). To correct for biases introduced by SNP density variations, the absolute allele-specific expression for mus and cas alleles was calculated by multiplying the bulk counts by the allelic ratio. Furthermore, for analysis involving X-reactivation, only genes biallelically expressed in iPSC and ESC were considered (allelic expression ratio >0.4 and <0.6), leaving 275 genes.

### Identification of A and B Compartments

Two Hi-C sub-matrices were extracted from normalized Hi-C matrices at 100-kb resolution with a split point at *Dxz4* (at coordinates 75.6 Mb). The A/B compartment scores were then computed separately with a standard principal-component approach. Namely, the matrix entries were normalized by the distance decay from the diagonal (*i.e.*, interaction scores were divided by the average interaction score at the given linear distance) and then transformed into correlation matrices using the Pearson product-moment correlation. The first principal component of a PCA (PC1) on each of these matrices was used as a quantitative measure of compartmentalization, and A+T-content was used to assign negative and positive PC1 categories to the correct compartments. When necessary, the sign of the PC1 (which is randomly assigned) was inverted so that positive PC1 values corresponded to the A compartment, and vice versa for the B compartment.

To more accurately define A/B-compartments, we then applied a gaussian mixture model with two components (k = 2) to the values of the PC1 using the R package mclust (Scrucca et al., 2016) using an unequal variance model (“V”). To reduce the impact of extreme outliers we used the 5% trimmed mean of values. The PC1 values were then centered at the intersection of the two gaussian models, separately for each genotype and sample. After centering, values were then normalized to +1 to -1.

To calculate the number of compartments, we separately merged bins of A and B compartments, positive and negative PC1 values respectively, using bedtools “merge” (Quinlan, 2014) and excluded bins smaller than 300 kb.

### Identification of Spatial Clusters and Subcompartments

Identification of spatial clusters and subcompartments was performed as described previously (Lucic et al., 2019). To smooth the diagonal decay and other biases, observed-over-expected normalization was applied individually to each contact matrix (Lieberman-Aiden et al., 2009), followed by ICE row-sum balancing (Imakaev et al., 2012). Outliers, such as enhancer-promoter loops, were clipped to the 90-th percentile value of each matrix. On the balanced matrix, the correlation matrix was computed and its diagonal was set to 0. The outliers of the correlation matrix were further smoothed with linear scaling; values below the 5 -th and above the 95-th percentiles were clipped to -1 and +1, respectively, and the values within this range were scaled proportionally.

Spatial clusters were identified on the correlation matrix running *k*-means (with 10 restarts) on *k*=12 weighted eigenvectors, i.e. the 12 leading eigenvectors, each weighted by its respective eigenvalue. Clustering of rows was only performed on D5 P-RFP+ X_mus_. Subcompartment names A1, A2, AB, B1, and B2 were assigned based on their mutual interaction patterns and their PC1 values in NPC X_mus_. For other time points, cluster labels were aligned with the ones found in D5 state.

### Insulation, TAD, and TAD Boundary Calling

Normalized contact matrices at 50-kb resolution were used to obtain the insulation score based on the number of contacts between bins on each side of a given bin, using a previously described method (Crane et al., 2015) with default parameters. To account for variation in SNP density, the signal of the insulation score was further smoothed by using moving averages of span 7 and selecting local minima as TAD borders. A cut-off value of -0.086 was chosen, as it gave a similar number of TADs on the active X as previously reported (Dixon et al., 2012), *i.e*., a TAD border was inserted at every local minimum with a smoothed insulation score below the cut-off.

### Domain Score

The domain score was used to quantify the degree of connectivity within TADs and was calculated as described previously (Krijger et al., 2016). Briefly, normalized contact matrices at 50-kb resolution were used. Then for each TAD, the fraction of intra-TAD contacts over its total number of *cis* contacts were used to calculate the domain score.

### Inter- and Intracompartment Strength Measurements

We followed a previously reported strategy to measure overall interaction strengths within and between A and B compartments (Schwarzer et al., 2017). Briefly, we based our analysis on the 100-kb bins showing the most extreme PC1 values, discretizing them by percentiles and taking the bottom 20% as the B compartment and the top 20% as the A compartment. We classified each bin in the genome according to PC1 percentiles and gathered contacts between each category, computing the log_2_ enrichment over the expected counts by distance decay. Finally, we summarized each type of interaction (A-A, B-B and A-B/B-A) by taking the median values of the log_2_ contact enrichment.

### Inter-Mega-Domain Interactions

To quantify the degree of inter-mega-domain interactions of subcompartments, we filtered depth-corrected Hi-C contact maps at 50-kb resolution for long-range chromatin interactions (>8.2 Mb) and then selected for strong interactions (top 20%). We then counted the number of strong long-range inter-mega-domain interactions and of all long-range inter-mega-domain interactions for each spatial cluster and then calculated the fraction of strong versus all long-range inter-mega-domain interactions.

### Integration of Published ChIP-Seq, CHART-Seq and DamID Data

Datasets generated by (Wang et al., 2018) were downloaded from GEO. BigWig files were converted to wig using bigWigToWig and then to bed using wig2bed (Neph et al., 2012). Lift over of bed files from mm9 to mm10 was performed using CrossMap (Zhao et al., 2014). Bedtools “map” and “intersect” functions were then used to integrate data with spatial clusters (Quinlan, 2014).

### Visualization

All plots with the following exceptions were generated using ggplot2 (Wickham, 2016). ATAC-Seq profiles and peaks, A/B compartments, and insulation score data were plotted using pyGenometracks (Ramírez et al., 2018). ATAC-Seq profiles for figure 5C were generated using GVIZ (Hahne and Ivanek, 2016). FACS plots were generated using FlowJo. ForceAtlas2 network of spatial cluster interactions was visualized using Gephi (Jacomy et al., 2014).

### Statistics and Reproducibility

RNA-Seq, ATAC-Seq and i*n situ* Hi-C data throughout the paper were generated by analysis of two biologically independent samples. Representative data are shown only if results were similar for both biologically independent replicates. All box plots depict the first and third quartiles as the lower and upper bounds of the box, with a band inside the box showing the median value and whiskers representing 1.5x the interquartile range. Wilcoxon rank-sum tests were performed with the wilcox.test() function in R in a two-sided manner.

### Resource Availability

Further information and requests for resources and reagents should be directed to and will be fulfilled by the Lead Contact, Bernhard Payer.

### Data and Code Accessibility

Data and analysis scripts will be released to public databases when the article is accepted for publication.

## Supplementary Information

**Figure S1.**
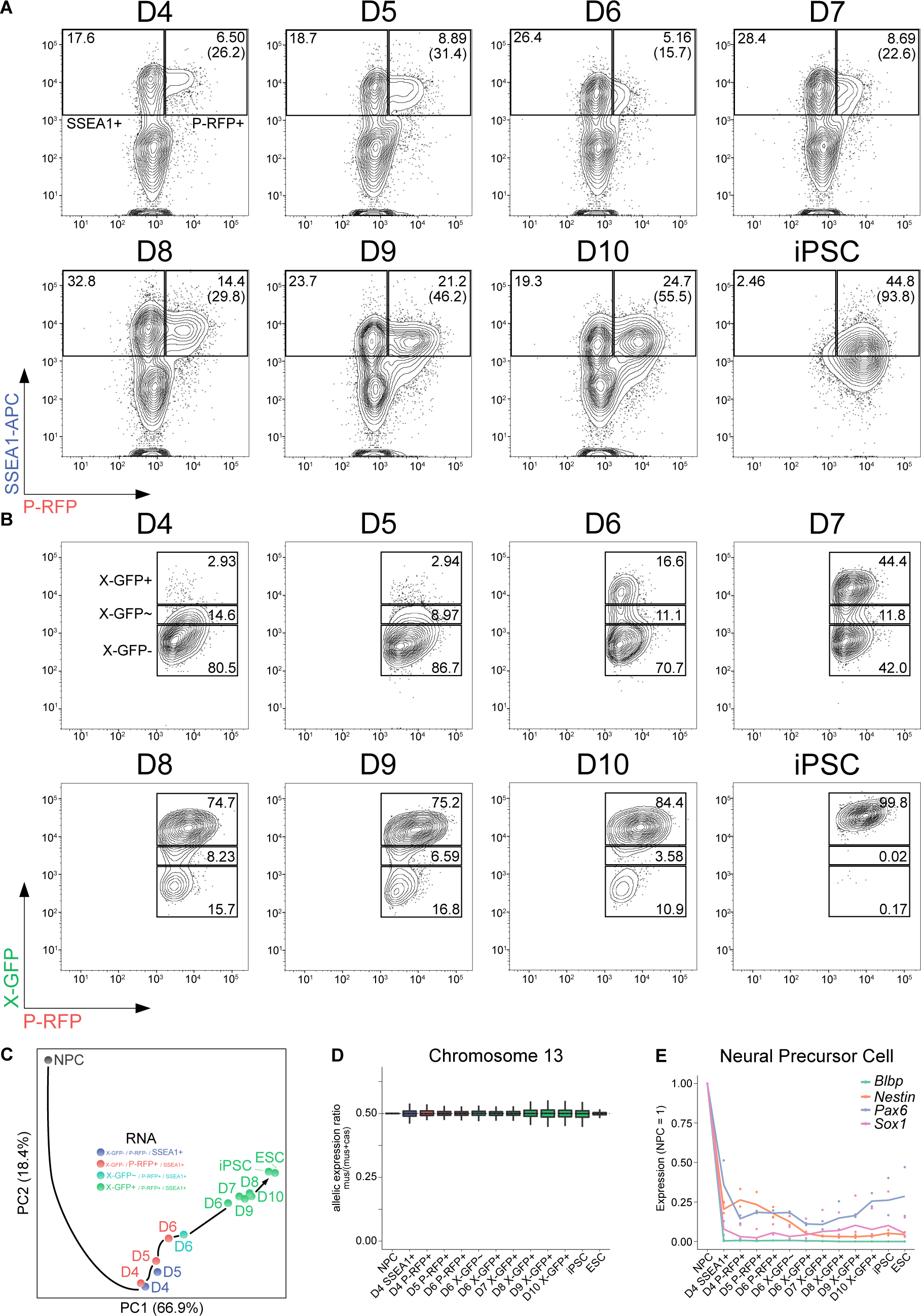
A Novel Reprogramming System to Efficiently Trace X-Chromosome Reactivation. (A) FACS gating strategy of P-RFP reporter expression during a reprogramming time course. Shown are representative contour plots gated on live cells. Numbers indicate the percentage of cells. Numbers in brackets indicate the percentage of P-RFP+ cells out of SSEA1+ cells. (B) FACS gating strategy of X-GFP reporter expression during a reprogramming time course. Shown are representative contour plots gated on SSEA1+/P-RFP+ cells in (A). Numbers indicate the percentage of cells. (C) PCA of dynamics of gene expression during reprogramming including neural precursor cells (NPCs) (n = 12,318 genes). Black arrow, hypothetical trajectory. (D) Allelic expression ratio (mus/(mus+cas)) of protein-coding genes expressed from chromosome 13 (n = 335). Biallelic expression, ratio = 0.5. (E) Average gene expression kinetics of neural precursor cell markers *Blbp*, *Nestin*, *Pax6*, and *Sox1* during reprogramming (n = 2, relative to the levels in NPC).

**Figure S2.**
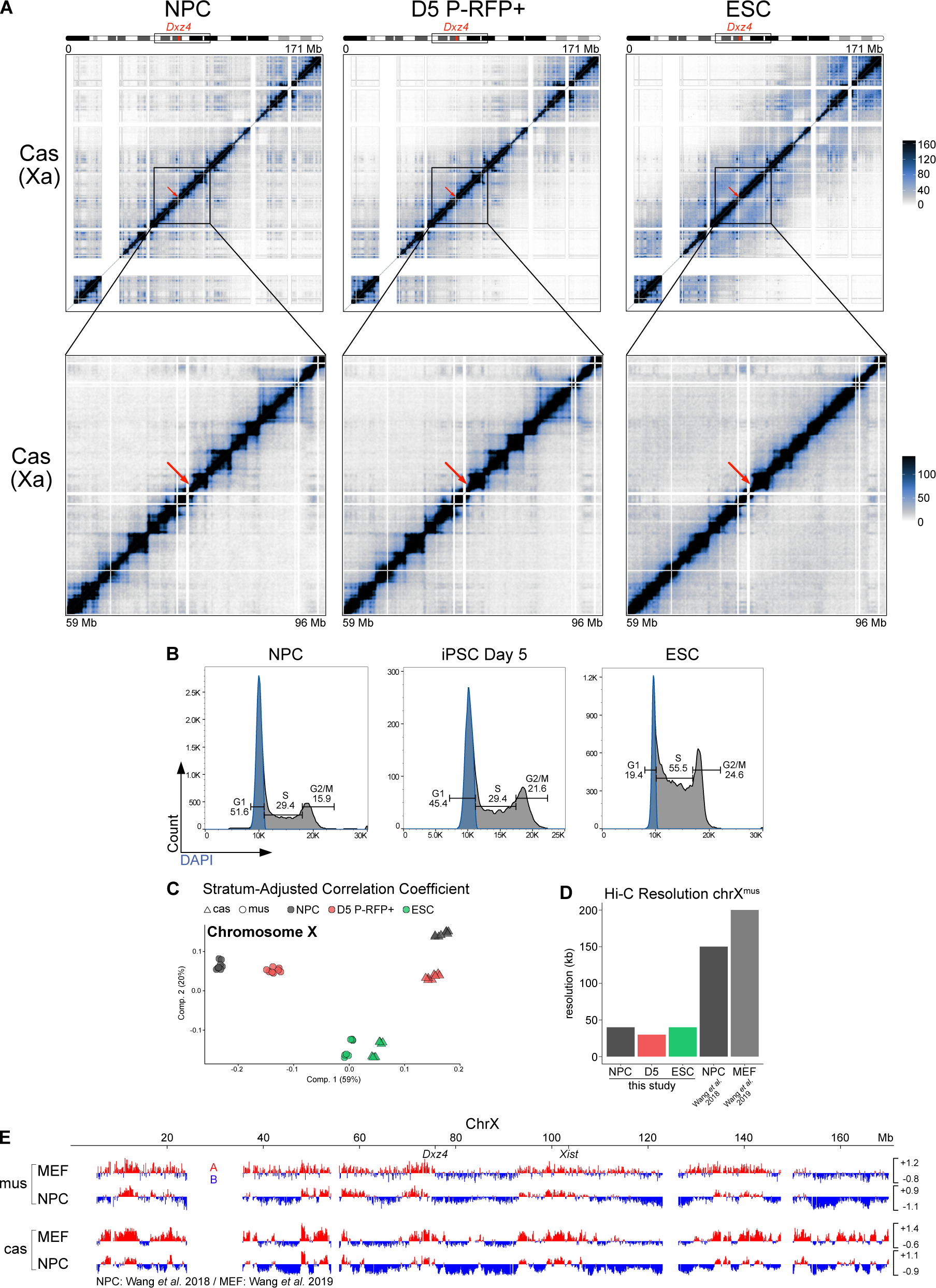
The Inactive X Chromosome Exhibits A/B-Like Compartmentalization. (A) Allele-specific Hi-C maps of the always active chromosome X^cas^ across stages. Top: Entire chromosome is shown at 200-kb resolution. Bottom: Zoom-in of the mega-domain boundary is shown at 100-kb resolution. Scale is shown in mega-bases (Mb). The mega-domain boundary *Dxz4* is indicated by a red arrow. White-shaded areas, unmappable regions. (B) FACS analysis of cell cycle using DAPI. Blue indicates sorted G1 population. Numbers indicate the percentage of cells. (C) Correlation of Hi-C samples at 100-kb resolution using stratum-adjusted correlation coefficient (Yang et al., 2017), visualized as a multi-dimensional scaling plot. (D) Hi-C resolution of the inactive chromosome X^mus^ achieved in this study, calculated as described previously (Rao et al., 2014). Resolution of comparable studies assessing the inactive X using allele-specific *in situ* Hi-C are shown (Wang et al., 2018, 2019). (E) A/B compartments of chromosome X at 100-kb resolution obtained with principal component analysis of matrices split at the *Dxz4* mega-domain boundary of neural precursor cells (NPCs) (Wang et al., 2018) and mouse embryonic fibroblasts (MEFs) (Wang et al., 2019). Positive PC1 values represent A-like compartments (red); negative PC1 values represent B-like compartments (blue).

**Figure S3.**
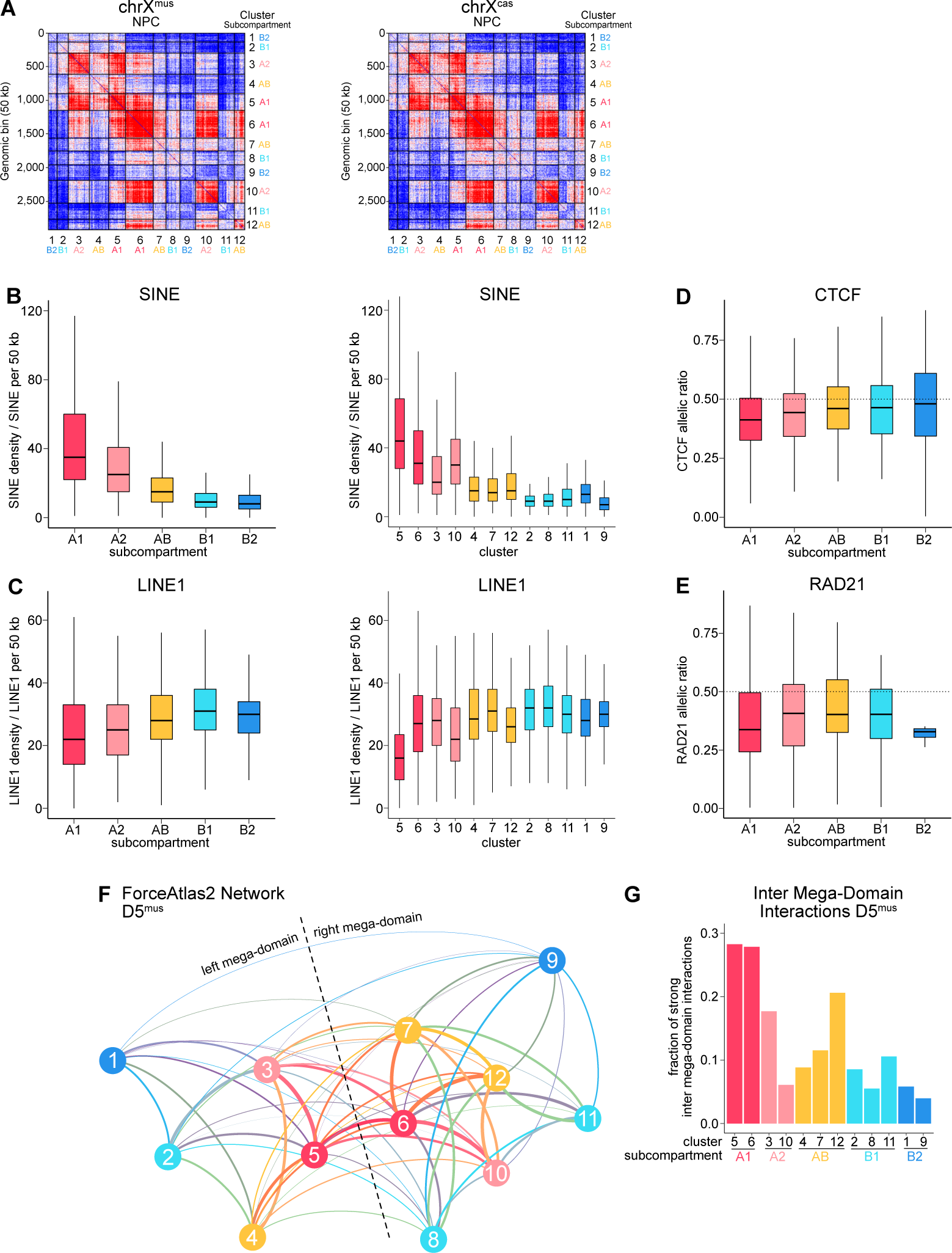
Subcompartmentalization of the Inactive X Chromosome. (A) Interaction matrices of spatial clusters at 50-kb resolution. (B) Density of SINE repeats per 50 kb bin for subcompartments and spatial clusters. (C) Density of LINE1 repeats per 50 kb bin for subcompartments and spatial clusters. (D) Allelic ratio of CTCF peaks between NPC^mus^ and NPC^cas^ of subcompartments. ChIP-Seq data from (Wang et al., 2018). (E) Allelic ratio of RAD21 peaks between NPC^mus^ and NPC^cas^ of subcompartments. ChIP-Seq data from (Wang et al., 2018). (F) Network of spatial clusters on chromosome X^mus^ in D5 P-RFP+ obtained by applying the ForceAtlas2 algorithm to Hi-C interaction patterns of spatial clusters. Each cluster represents a single node of the network. Line-width correlates with interaction strength. (G) Inter-mega-domain interactions of clusters (across the mega-domain boundary) in D5 P-RFP+^mus^.

**Figure S4.**
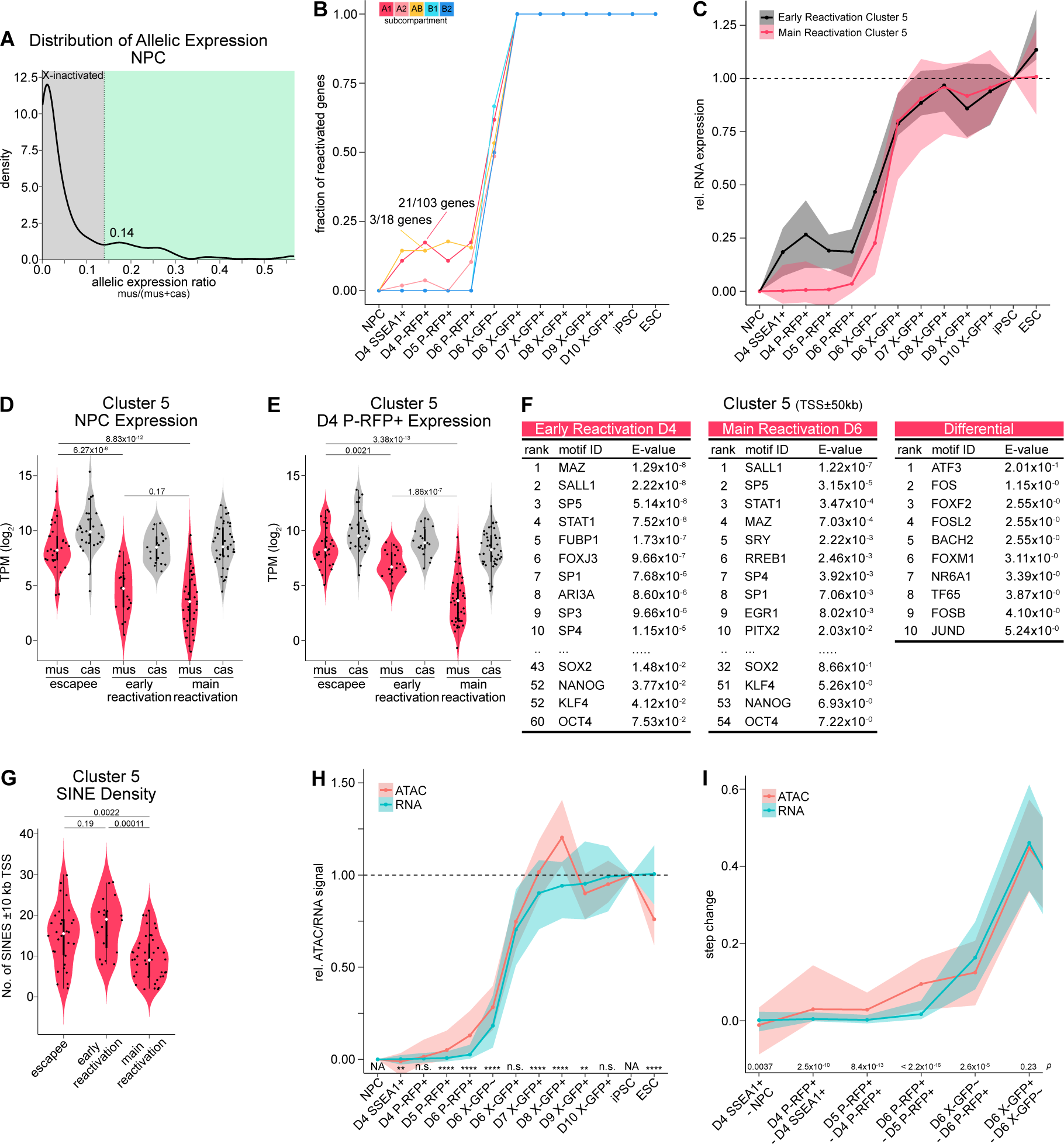
Initiation of Chromatin Opening and Gene Expression from a Distinct 3D Cluster. (A) Distribution of the allelic expression ratio of chromosome X in NPCs (n = 275). The dashed line represents the cut-off of 0.14, above which genes were considered biallelically expressed (green shading). Only protein-coding genes with sufficient allelic information, expression for chromosome X^cas^, and biallelic expression in iPSC and ESC are counted (*see methods*). (B) Dynamics of gene reactivation of subcompartments. Fractions of reactivated genes per cluster are shown. 0, no reactivated gene. 1, all genes reactivated. Threshold for gene reactivation, allelic expression ratio >0.14 (see A). While subcompartment AB shows a similar fraction of early reactivating genes to A1, it is biased by the low number of genes in this subcompartment, with a total of 18 genes and 3 reactivating at D4 P-RFP+. (C) Relative RNA expression (NPC = 0, iPSC = 1) for genes of cluster 5 reactivating early, at D4 P-RFP+, compared to genes reactivating after that (“main reactivation”). Line shows median. Shading denotes 50% confidence interval. (D) Violin plots showing the RNA expression from the mus allele of genes of cluster 5 in NPCs. TPM, transcripts per million. The p values are calculated by Wilcoxon rank-sum test. (E) As (D) in D4 P-RFP+ cells. (F) Enrichment of transcription factor motifs. AME was used to identify differentially enriched motifs from ATAC-Seq peaks in a window of +-50 kb around the TSS of cluster 5 genes reactivating early or after that. Differential AME analysis for early reactivation-specific motifs was performed using peaks around early reactivating genes as primary sequence and peaks around main reactivating genes as control. (G) Violin plots showing the number of SINE repeats in a window of +-10 kb around the TSS of genes of cluster 5. The p values are calculated by Wilcoxon rank-sum test. (H) Relative gene expression and chromatin opening compared (NPC = 0, iPSC = 1). Line shows median. Shading denotes 50% confidence interval. The p values are calculated by Wilcoxon rank-sum test. (I) Step changes of gene expression and chromatin opening compared. The p values are calculated by Wilcoxon rank-sum test.

**Figure S5.**
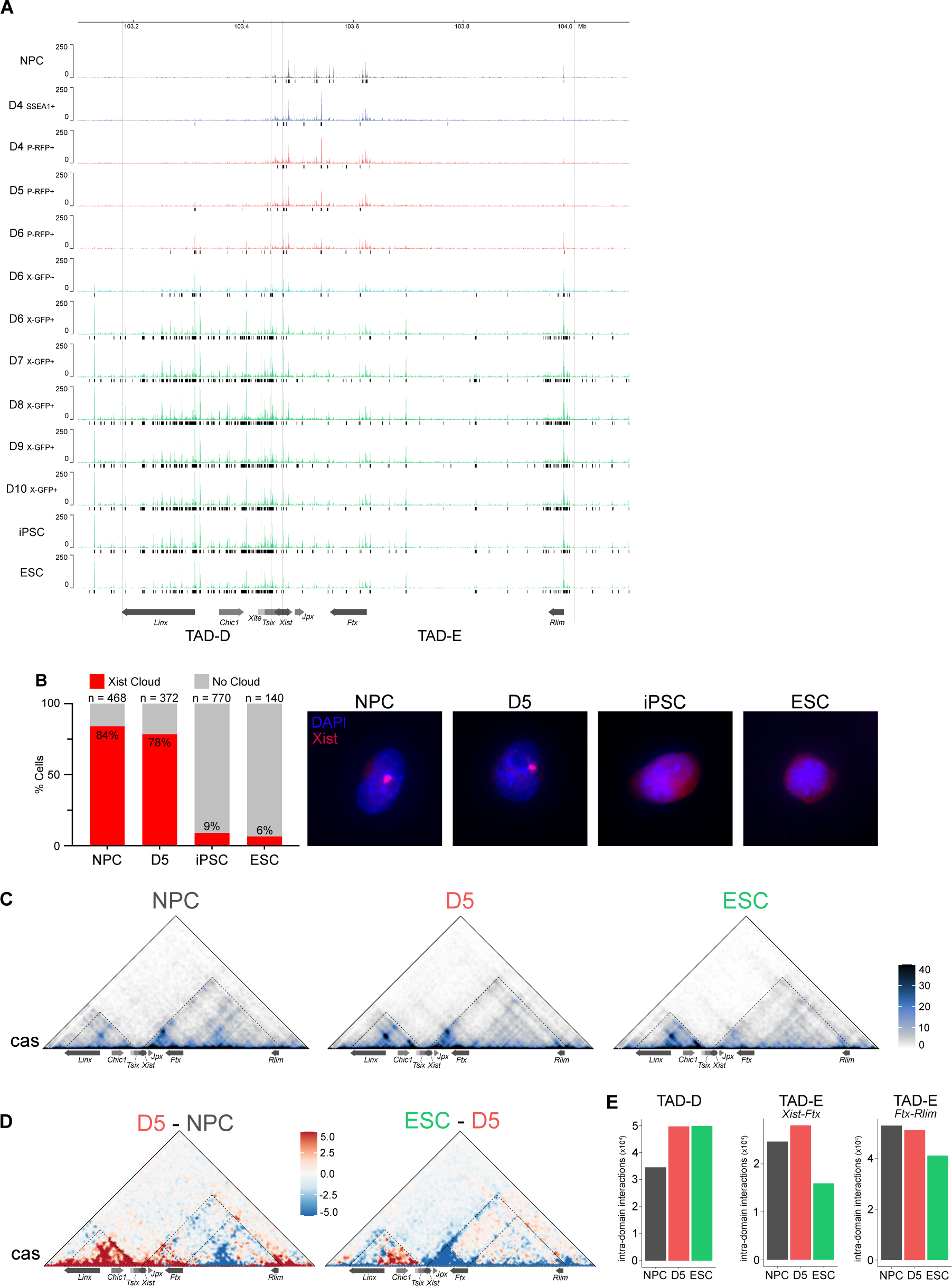
Remodeling of the X-inactivation Centre Leading to *Xist* Downregulation. (A) ATAC-Seq profiles at a region encompassing *Tsix* TAD-D (mm10; 103.18 Mb - 103.45 Mb) and *Xist* TAD-E (mm10; 103.47 Mb - 104.0 Mb). ATAC-peaks in black (except for NPCs, differential peaks compared to NPCs are shown. Only genes with implicated roles in X-inactivation or X-reactivation are shown. (B) Downregulation of *Xist* RNA during reprogramming based on RNA-FISH. Percentages of cells containing *Xist* RNA FISH clouds are shown. (C) Allele-specific Hi-C maps of chromosome X^cas^ at 10-kb resolution at a region encompassing TAD-D (left) and TAD-E (right). Dotted lines show TADs. (D) Differential allele-specific Hi-C maps of chromosome X^cas^ at 10-kb resolution at a region encompassing TAD-D and TAD-E. Dotted lines show TADs and additionally separates TAD-E in two for quantification in (**E**). (E) Sum of intra-domain interactions are shown. TAD-E was separated in two regions at the TSS of *Ftx*.

**Figure S6.**
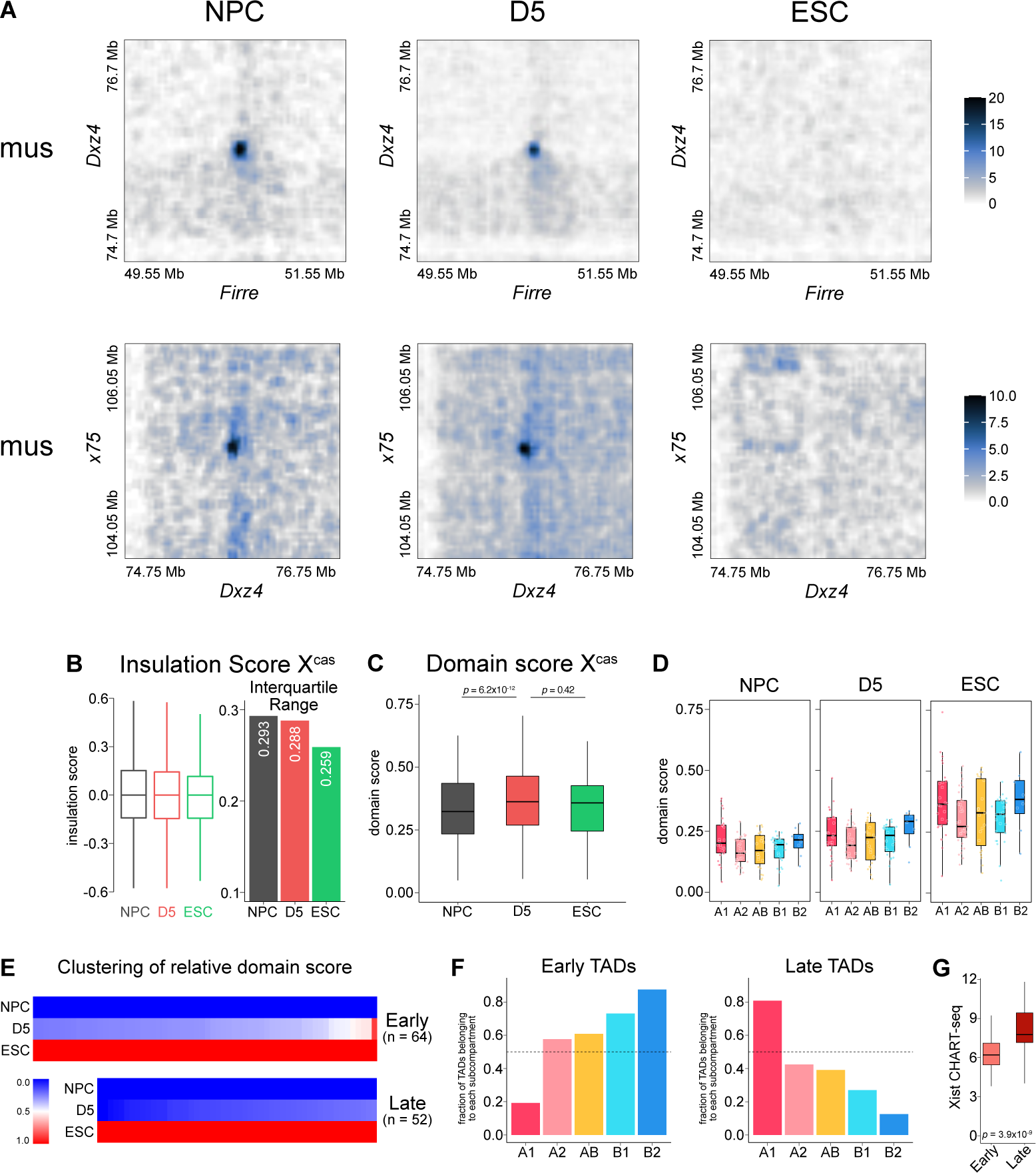
Structural Changes During X-Reactivation in the Absence of Chromatin Opening and Transcription. (A) Allele-specific Hi-C maps at 30-kb resolution showing superloop interactions between *Dxz4* and *Firre*, as well as *x75* and *Dxz4* ± 1 Mb. (B) Comparison of insulation scores for chromosome X^cas^. Interquartile range of insulation scores is shown on the right. (C) Comparison of domain scores for chromosome X^cas^. The p values are calculated by Wilcoxon rank-sum test. (D) Domain scores for X^mus^ of subcompartments. (E) *k*-means clustering (k = 2) of relative domain scores to identify early and late TADs. (F) Comparison of fraction of TADs of each subcompartment belonging to “early” or “late” TADs. (G) Xist RNA enrichment of early and late TADs in NPCs (composite scaled data). CHART-Seq data from (Wang et al., 2018).

